# Unraveling the Interplay between Stability and Flexibility in Design of Polyethylene Terephthalate (PET) Hydrolases

**DOI:** 10.1101/2024.05.11.593663

**Authors:** Shiqinrui Xu, Chengze Huo, Xiakun Chu

## Abstract

The accumulation of polyethylene terephthalate (PET), a widely used polyester plastic in packaging and textiles, poses a global environmental crisis. Biodegradation presents a promising strategy for PET recycling, with PET hydrolases (PETase) undertaking the task at the molecular level. Unfortunately, due to its low thermostability, PETase can only operate at ambient temperatures with low PET depolymerization efficiency, hindering its practical application in industry. Currently, efforts to engineer PETase have primarily focused on enhancing its thermostability. However, increased stability often reduces the structural dynamics necessary for substrate binding, potentially slowing down the enzymatic activity. To elucidate the delicate balance between stability and flexibility in optimizing PETase catalytic activity, we performed theoretical investigations on both wild-type PETase (WT-PETase) and a thermophilic variant (Thermo-PETase) using molecular dynamics simulations and frustration analysis. Despite being initially designed to stabilize the native structure of enzyme, our findings reveal that Thermo-PETase exhibits an unprecedented increase in structural flexibility at the PET binding and catalytic sites, beneficial for substrate recruitment and product release, compared to WT-PETase. Upon PET binding, we observed that structural dynamics of Thermo-PETase are largely quenched, facilitating subsequent chemical reactions. Compared to WT-PETase, Thermo-PETase forms more extensive interactions with PET, resulting in a higher population of catalytically competent enzyme-substrate states, thus contributing to increased catalytic activity. Our theoretical results are consistent with experimental findings and further suggest that Thermo-PETase exhibits higher catalytic activity than WT-PETase across a broad temperature range by leveraging stability and flexibility at high and low temperatures, respectively. Our findings offer valuable insights into how PETase optimizes its enzymatic performance by balancing stability and flexibility, paving the way for future PETase design strategies.

## 1 Introduction

Plastics play a vital role in all aspects of everyday life, offering numerous benefits to the development of modern society. However, due to the ultralong lifetimes of most synthetic plastics, plastic waste accumulation has now become one of the most globally challenging environmental crises, profoundly impacting the ecosystems and posing great health risks to both wildlife and humans [1, 2, 3]. Among the various types of plastics, polyethylene terephthalate (PET) is the most abundant thermoplastic polyester manufactured in the world, primarily because of its widespread usage in the packaging market and textile industry [4, 5]. Despite its prevalence, PET exhibits high resistance to degradation, leading to the accumulation of PET plastic waste after use and contributing significantly to the solid waste problem worldwide [6, 7].

Great efforts have been made towards PET biodegradation, which is an environmentally friendly and efficient technology, ideal for the closed-loop recycling of PET plastics [8]. Enzymes that exhibit PET-degradation activity are known as PET hydrolases, which are capable of breaking down the longchain PET molecules into their building blocks [9, 10, 11]. Over recent decades, several PET hydrolases have been identified and studied, including cutinases, lipases and esterases [12, 13]. Among these, PET-degrading cutinases from thermophilic micro-organisms have shown remarkable PET degradation efficiency at high temperatures, near the glass transition temperature of PET, where the PET crystalline structure starts to melt towards the amorphous polymer chains [13, 14, 15, 16, 17]. In 2016, Yoshida et al. made a breakthrough discovery of a cutinase-like PET hydrolase, named PETase, in the bacterium *Ideonella sakaiensis*, which can degrade and assimilate PET as its source of carbon and energy [18]. PETase is a naturally evolved PET hydrolase, exhibiting high PET depolymerization activity at ambient temperatures. Consequently, this unique property of PETase has distinguished it from PET-degrading cutinases [11]. However, PETase showed very low degradation efficiency for highly crystallized PET and rapidly lost its enzymatic activity with increasing temperature [19]. These features impede the practical applications of PETase in the plastic degradation industry.

To increase the thermostability of PETase and further improve the PET degradation performance, various engineering strategies have been devised [20,21,22,23,24,25,26,27,28,29,30, 31]. Son et al. discovered that PETase with only three-residue substitution (S121E/D186H/R280A) remarkably enhanced the PET depolymerization activity across different ambient temperatures, compared to wild-type PETase (WT-PETase) [21]. This new variant, named Thermo-PETase, derived from a structure-based protein engineering strategy, was primarily designed to stabilize the flexible loop, which exhibited high B-factor values in the crystal structure. The mutations proposed for Thermo-PETase established the newly formed interactions in the native structure of PETase, thus increasing the melting temperature by 8.8K compared to WT-PETase [21]. Similar strategies focusing on mutations of residues to stabilize the most flexible loop in PETase, have been recently developed to successfully achieve the increased thermostability [32, 33]. Moving forward, a machine-learning method based on training with nearly 19,000 structures from the Protein Data Bank (PDB) was employed to introduce two additional mutations to Thermo-PETase, giving rise to FAST-PETase [25]. FASTPETase, which demonstrated much higher PET degradation activity than many other PETase variants, exhibited an increase in melting temperature by 9K over its scaffold Thermo-PETase. Structural analysis reveals that the increased thermostability of FAST-PETase is largely attributed to the formation of favorable salt bridges and hydrogen bonds in the native structure. Recently, directed evolution methods have been applied to PETase design, resulting in successful variants capable of operating at the glass transition temperature of PET, presenting a significant step towards the complete PET depolymerization [26, 29].

Despite significant achievements made in the discovery of highly thermostable PETase variants [34, 35, 36], progress towards finding PETase capable of operating efficiently at ambient temperatures has been proceeding slowly. This is largely due to the fact that our understanding of the structure-function relationship in PETase remains elusive [37, 19, 38, 39]. It has been recognized that, unlike other thermophilic cutinases, the unique property of PET degradation activity at ambient temperatures for PETase is attributed to its open and flexible substrate binding cleft [40]. Increasing evidence has revealed that modulating the opening of the binding cleft in thermophilic cutinases can enhance their catalytic efficiency on PET degradation [41, 42], highlighting the importance of structural flexibility at the substrate binding site of PET hydrolases in facilitating enzymatic activity. However, by narrowing down the binding cleft on the surface of WT-PETase through mutations, Austin et al. observed increased hydrolytic activity on PET [38], leading to the seemingly conflicting effects of structural dynamics in PETase on exerting its catalytic function. Furthermore, increasing the thermostability of PETase is generally achieved by introducing newly formed interactions to stabilize the native structure [43, 44]. Thus, reducing the structural flexibility at the native state of PETase, a common strategy for improving stability in PETase design, may be disadvantageous to substrate binding and product release, potentially slowing down the catalytic activity. Understanding the interplay between stability and flexibility in PETase is fundamental and crucial for further engineering efficient PET hydrolase operating at ambient temperature, but it presents great challenges.

In this work, we performed molecular dynamics (MD) simulations along with frustration analysis on WT-PETase and one prototypical thermophilic PET variant, Thermo-PETase, which was observed to exhibit an overall enhanced PET hydrolytic activity over WT-PETase across a wide range of temperatures [21, 25]. Consistent with experimental findings, our simulations show that Thermo-PETase unfolds more slowly than WT-PETase at high temperature [21], thus allowing ThermoPETase to maintain its functional activity for a longer duration at elevated temperatures. Interestingly, despite the increase in global stability led by mutations, we observed that ThermoPETase possesses more significant structural flexibility than WT-PETase at ambient temperatures. Our detailed analyses unveil that the structural dynamics at the PET binding site and catalytic triad in Thermo-PETase are more pronounced, potentially facilitating substrate recruitment and product release, compared to WT-PETase. Additionally, frustration results indicate that there are large-scale arrangements of frustrated contacts in Thermo-PETase upon mutations, leading to an increased degree of frustration at the local binding site while maintaining global scales unchanged. Moreover, we found that the structural dynamics of Thermo-PETase within the substrate-enzyme complex are largely quenched by the interactions formed with PET, resulting in a stable catalytic state primed for subsequent highly efficient chemical reactions. Our results provide valuable insights into the future rational design of PETase and other PET hydrolases towards PET degradation at ambient temperature by balancing the global stability and local flexibility.

## 2 Results

### 2.1 Structural dynamics of PETase at native states

We performed microsecond-long molecular dynamics (MD) simulations initialized from the native structures of WT- and Thermo-PETase to investigate the temperature-dependent structural dynamics of PETase. MD simulations were carried out at two ambient temperatures (298K and 308K) and one elevated temperature (450K), which is much higher than the melting temperatures of WT- and Thermo-PETase measured by experiments (321.8K and 329.8K) [21]. Root mean square deviation (RMSD) relative to the respective native structures was then calculated for each simulation. We observed that Thermo-PETase consistently exhibits lower RMSD values than WT-PETase at all three temperatures, in particular at the high temperature of 450K (Figure 1A-C, Figure S2). These results suggest that Thermo-PETase possesses enhanced thermodynamic stability upon the mutations, consistent with experimental observations [21]. Additionally, the consistently lower RMSD values of Thermo-PETase compared to WT-PETase at both 298K and 308K imply an overall reduction in the structural dynamics of Thermo-PETase relative to WT-PETase.

**Figure 1:**
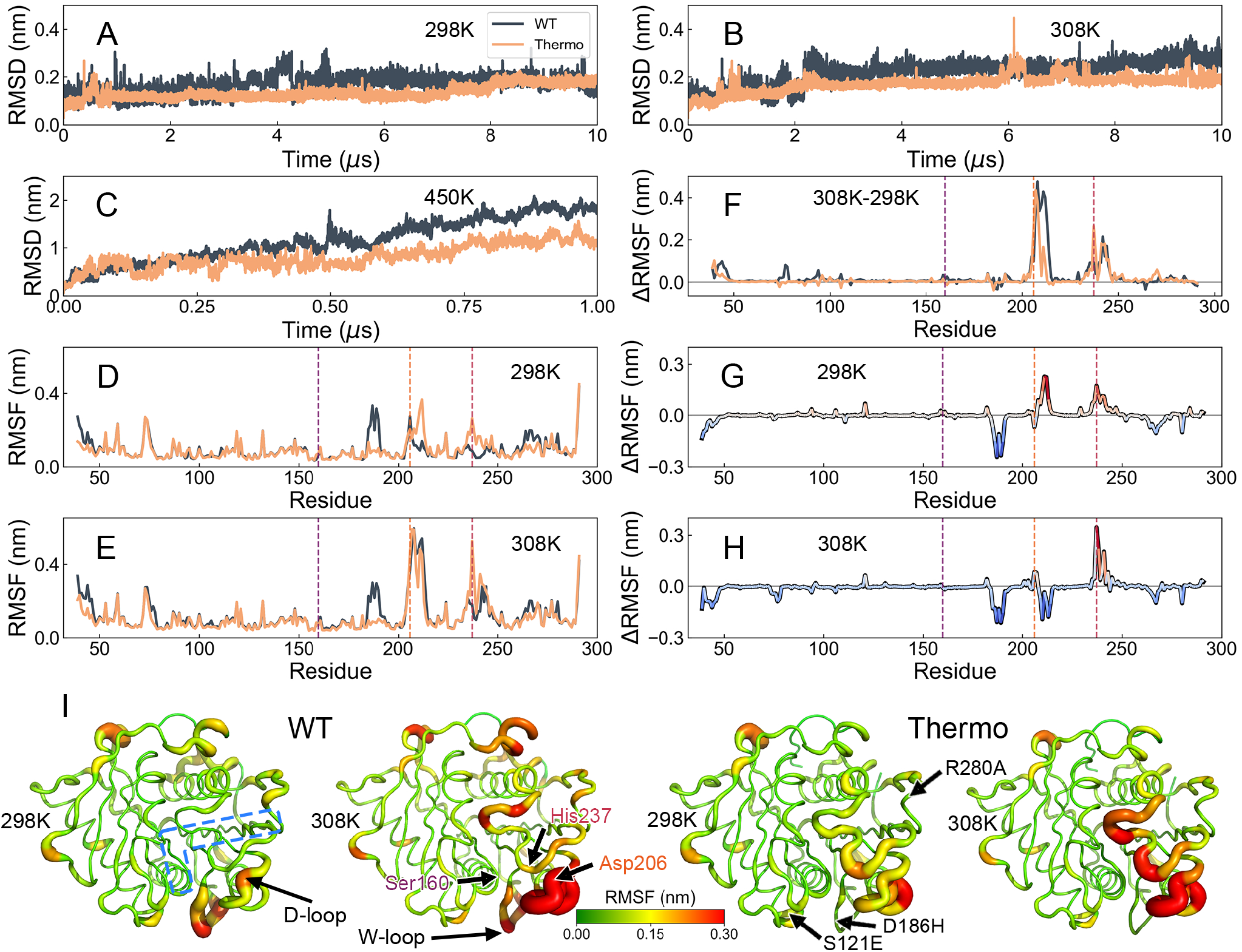
Structural stability and flexibility of WT- and Thermo-PETase revealed by molecular dynamics (MD) simulations. Root Mean Square Deviation (RMSD) relative to the corresponding native structures of WT- and Thermo-PETase (PDB: 5XJH [19] and 6IJ6 [21]) for the simulations performed at (A) 298K, (B) 308K and (C) 450K, respectively. Root Mean Square fluctuation (RMSF) of WT- and Thermo-PETase for the simulations performed at (D) 298K and (E) 308K. (F) RMSF differences between 298K and 308K for WT- and Thermo-PETase. RMSF differences between WT- and Thermo-PETase (RMSF_*Thermo*_ RMSF_*WT*_) at (G) 298K and (H) 308K. The purple, orange, and red vertical lines in (D-H) indicate the catalytic triad Ser160, Asp206, and His237, respectively. (I) RMSF values at 298K and 308K for WT- and ThermoPETase projected onto the corresponding native structures. The thickness and color of the tubes illustrated on the PETase structures denote the magnitudes of RMSF. The blue dashed region in (I) illustrates the PET binding site on the surface of PETase. The PET binding site is determined by the docking of one PET tetramer to the WT-PETase, described in Figure S1. The three mutations (S121E/D186H/R280A) in Thermo-PETase with respect to WT-PETase are indicated in (I) [21].

To quantitatively assess the structural dynamics of PETase at the residue level, we calculated root mean square fluctuation (RMSF) by collecting all the simulation trajectories each at 298K and 308K (Figure 1D and 1E). The RMSF profiles and residue-level contact probability maps of WT- and Thermo-PETase show substantial overlap at both correspond-ing temperatures, with notable differences primarily observed at the region involving Trp185 (within the W-loop, Ala180– Pro197), and the region involving Asp206 (within the D-loop, Glu204–Ser213) and His237 of the catalytic triad (Figure S3). As temperature increases, the structural flexibility of these two catalytic residues (Asp206 and His237) in both WT- and Thermo-PETase appears to be significantly enhanced (Figure 1F). Interestingly, although the overall structural dynamics of WT-PETase is more significant than those of Thermo-PETase, as indicated by RMSD (Figure 1A), Thermo-PETase exhibits more flexibility in regions involving two catalytic residues at 298K (Figure 1G). Upon increasing the temperature to 308K, the D-loop in Thermo-PETase appears to be more stable than it in WT-PETase, while the flexibility of the region close to His237 in Thermo-PETase is still more pronounced than that in WT-PETase, as demonstrated by RMSF (Figure 1H) and contact probability map (Figure S3). The W-loop, which was identified as one of the most flexible regions in WT-PETase, has negative effects on its overall thermostability [45, 32]. Mutation on the residues within the W-loop (e.g., Asp186) in order to form stable electrostatic interactions or hydrogen bonds has been demonstrated as a practice strategy for enhancing the thermostability of PETase [21, 33]. Our simulations reveal that mutations in Thermo-PETase stabilize the W-loop by reducing the structural flexibility, potentially contributing to increased stability. On the other hand, the W-loop in Thermo-PETase remains consistently more stable than it in WT-PETase across different temperatures. This observation is reminiscent of the findings from a recent study, which has underlined the advantageous role of a rigid, correctly oriented Trp185 in stabilizing the binding interactions with the substrate [33].

To better illustrate the structural flexibility of PETase and its changes with increasing temperature, we mapped the RMSF values onto the native structures of WT- and Thermo-PETase (Figure 1I). Noteworthy, the RMSD between the PDB structures of WT- and Thermo-PETase is only 0.2Å, indicating that these two PETases share a high degree of similarity in the static native structure. However, notable differences in structural dynamics are evident, particularly in regions close to the PET binding site. Furthermore, at the low temperature of 298K, the Dand W-loops in WT-PETase exhibit certain degrees of flexibility, while most regions in WT-PETase are relatively static, including the PET binding site. As the temperature increases to 308K, the binding site of WT-PETase becomes dynamic. The flexible binding site in PETase may potentially facilitate substrate binding, contributing to the increased PET degradation efficiency [40]. In contrast, despite having a more rigid W-loop, Thermo-PETase at 298K exhibits excessive structural flexibility at the binding site, which becomes even more dynamic as the temperature increases to 308K. These distinct observations of structural dynamics between WT- and ThermoPETase may influence substrate binding, thus further making distinct contributions to the catalytic activity of PETase.

### 2.2 Free energy landscapes of PETase at native states

In order to capture the essential structural dynamics of PETase at the native states, we collected the simulation trajectories of WT- and Thermo-PETase at the same ambient temperatures (298K and 308K), where both PETases remained unfolded, and subsequently performed principal components analysis (PCA). At 298K, the free energy landscapes (FELs) projected onto the first and second principal components (PC1 and PC2) reveal that WT-PETase explores a notably smaller conformational space compared to Thermo-PETase, indicating significant structural flexibility in Thermo-PETase at low temperature (Figure 2A and 2B). At 308K, although the area of FELs is comparable for WT- and Thermo-PETase, the conformational spaces explored by these two PETases are significantly different, leading to distinct structural dynamics (Figure 2C and 2D). Further projections of simulation trajectories onto the PCs with structural illustrations showcase the key motions in PETase (Figure 2E-H). We observed that the most dominant structural dynamics in these two PETases at both temperatures are related to the collective motions led by the PET binding site and the catalytic site. Specifically, at 298K, the D-loop exhibits a twisting motion along PC1, whereas it swings in the opposite direction to the loop hosting His237 along PC2. This motion along PC2 may correspond to a conventional “open-to-close” conformational transition in PETase [38, 46], potentially in favor of accommodating PET binding. Furthermore, the FEL of WT-PETase features a single, wide basin, which includes the native state (denoted as state 1) and a native-like state 2. In contrast, the FEL of Thermo-PETase displays three widely distributed energy basins, with the most probable one corresponding to the native state. Thermo-PETase within the right most basin (denoted as state 4) and the intermediate basin (denoted as state 3) both show a more elongated D-loop (Figure 2I), which may facilitate PET binding [47].

**Figure 2:**
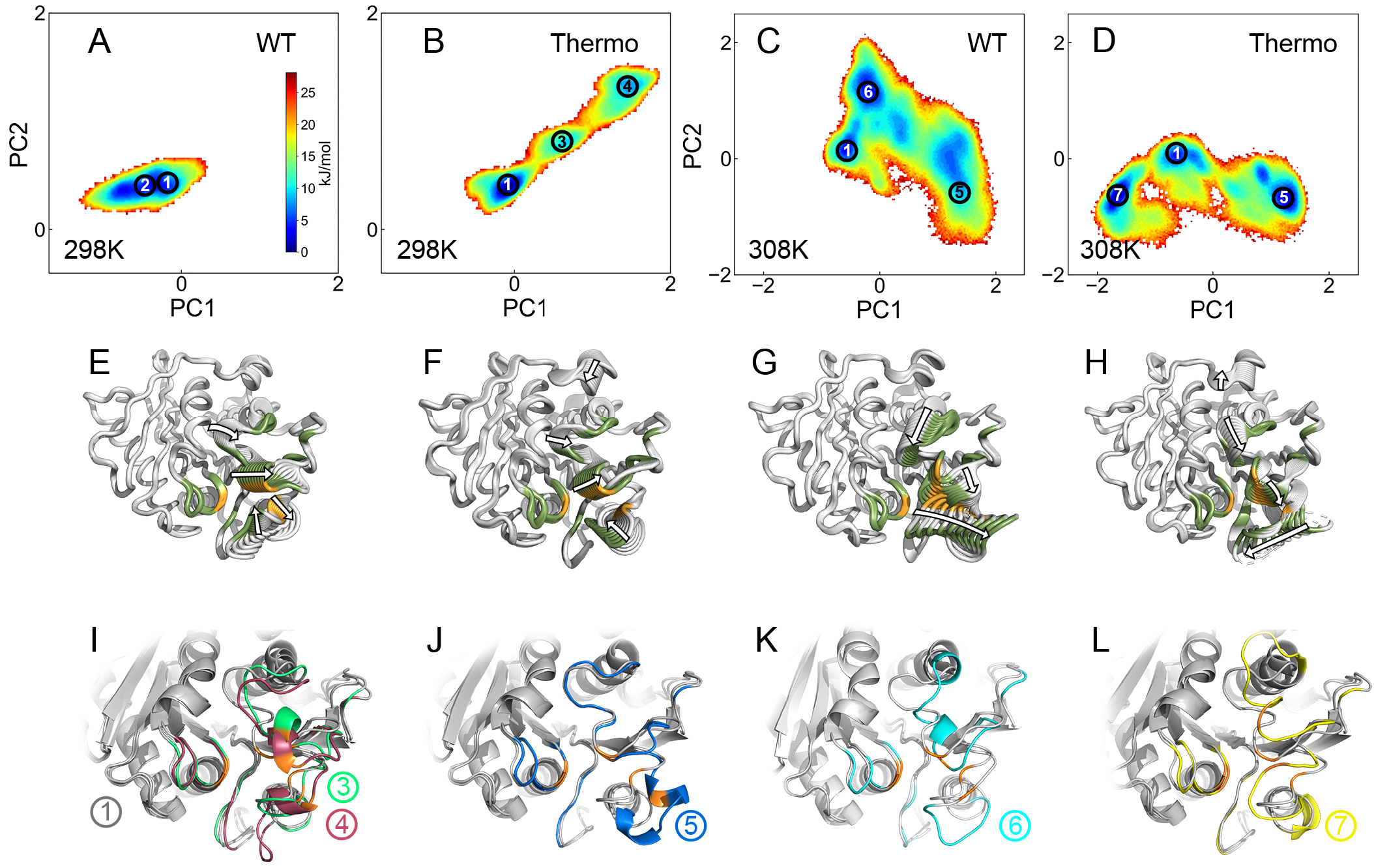
Principal component analysis (PCA) projected on the native dynamics of WT- and Thermo-PETase. (A, B) Free energy landscapes (FELs) of WT- and Thermo-PETase at 298K, projected onto the first and second principal component (PC1 and PC2). The weights of PC1 and PC2 are 20.89% and 15.35%, respectively. (C, D) Free energy landscapes (FELs) of WT- and Thermo-PETase at 308K, projected onto the PC1 and PC2. The weights of PC1 and PC2 are 29.19% and 12.18%, respectively. Representative structures of PETase extracted from the simulation trajectories at 298K projected onto (E) PC1 and (F) PC2, respectively. Representative structures of PETase extracted from the simulation trajectories at 308K projected onto (G) PC1 and (H) PC2, respectively. In each panel of (E-H), 10 representative structures ranging from the lowest to highest PC values are shown, with arrows indicating the structural motions along the corresponding PC. (I-L) Local PETase structural illustrations focusing on the PET binding site for different (meta)stable states indicated in (A-D). The PDB structure of WT-PETase (grey, representing state 1) is aligned with the (I) states 3 (green) and 4 (red), (J) state 5 (blue), (K) state 6 (cyan) and (L) state 7 (yellow). In (E-H), the residues at the PET binding site are colored in dark green. In (E-L), the three catalytic residues (Ser160, Asp206 and His237) are colored in orange. State 2 is structurally similar to state 1 (native structure), thus it is not shown. Full PETase structural illustrations are shown in Figure S4.

The collective motion patterns in PETase vary significantly as temperature changes. When the temperature increases to 308K, the structural dynamics of PETase along PC1 and PC2 predominantly concentrate on the PET binding site and are strongly related to the “open-to-close” transition (Figure 2G and 2H). In this respect, state 5, which was found to be highly populated in both WT- and Thermo-PETase, exhibits an open binding site (Figure 2J), serving as a binding-competent state. In addition, distinct structural dynamic behaviors were observed for these two PETase, as evidenced by the more widely distributed FEL of WT- and Thermo-PETase focusing along PC2 and PC1, respectively. It is worth noting that the collective motion of PETase along PC2 involves a twisting mode, which may lead to an incomplete opening of the binding site for PET. Careful analysis on the structures of PETase in states 6 and 7 (Figure 2K and 2L), which are respectively populated by WT- and Thermo-PETase, reveals no clear expansion in the binding cleft, thus their roles in PET binding remains elusive.

To elucidate the structural dynamics of PETase at the catalytic site, we projected the FELs onto the spatial distances between every pair of residues in the catalytic triad of Ser160, Asp206, and His237 (Figure 3). At 298K (Figure 3A-F), WT-PETase explores a much narrower conformational space than Thermo-PETase, with two (meta)stable states observed (states 1 and 2). Noteworthy, while WT-PETase in state 2 is overall structurally similar to the native structure shown in Figure 2, it possesses a slightly opened catalytic triad, compared to that in state 1. The existence of state 2 for WT-PETase may facilitate this enzyme to bind and consequently degrade PET at room temperature [40]. On the other hand, Thermo-PETase in states 3 and 4 exhibits similarly increased pairwise distances among these three catalytic residues compared to WT-PETase. This indicates that the structural flexibility of Thermo-PETase at the catalytic site is significantly enhanced by the mutations, thus promoting the binding of PET.

**Figure 3:**
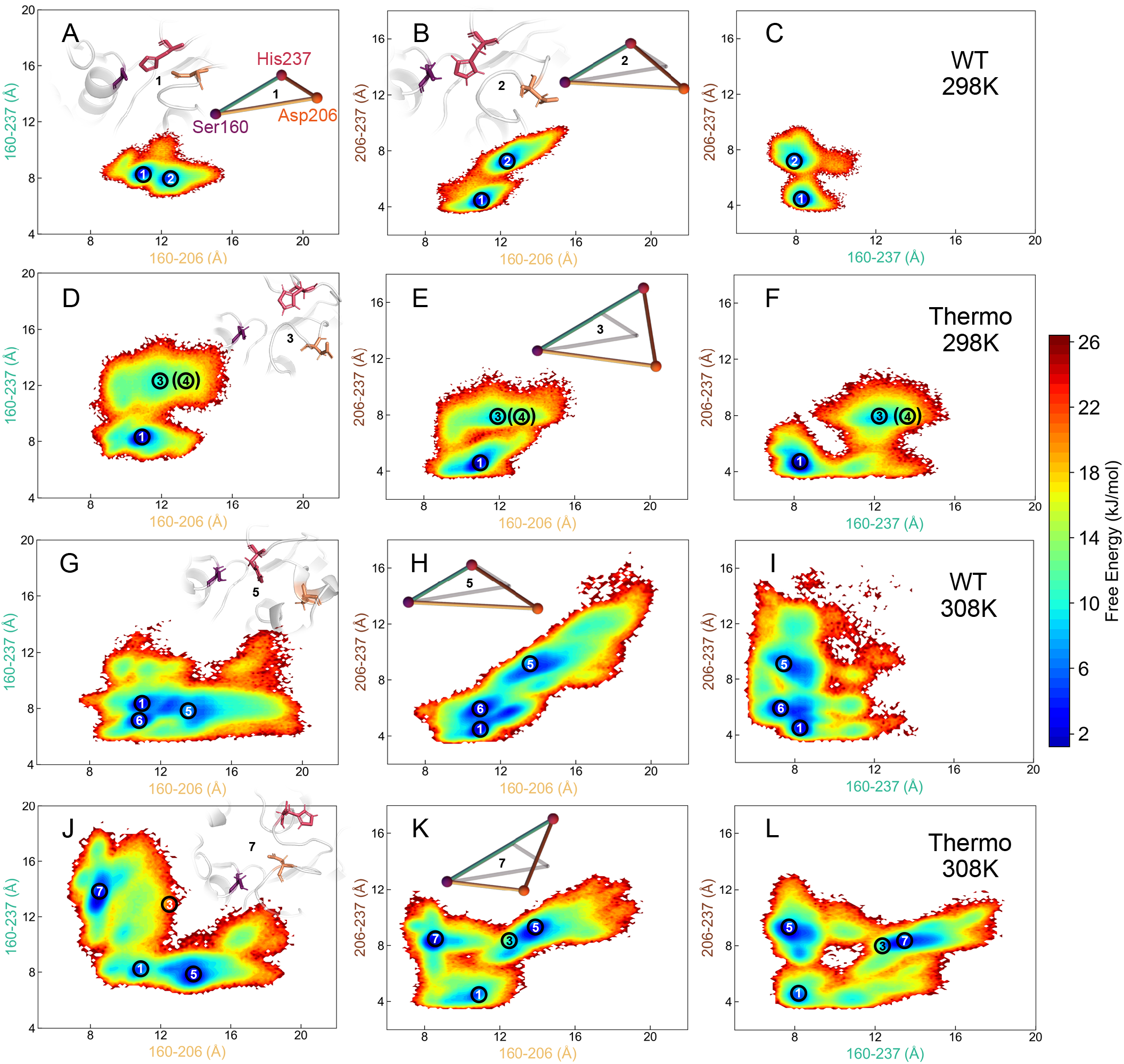
FELs projected onto the distances between every pair of the catalytic triad of Ser160, Asp206 and His237. FELs at 298K for (A-C) WT-PETase and (D-F) Thermo-PETase. FELs at 308K for (G-I) WT-PETase and (J-L) Thermo-PETase. Typical structures of PETase focusing on the catalytic triad at different (meta)stable states are shown. The (meta)stable states are numbered in the same was as they are in Figure 2. Colored triangles depict the geometries of the catalytic sites of PETase at different (meta)stable states, while the grey triangle depicts the geometry of the catalytic site in the native structure of WT-PETase (state 1). The pairwise distances of the catalytic residues in state 6 are similar to those in state 1 (native structure), thus the structural illustration of state 6 is not shown.

At 308K (Figure 3G-L), although WT- and Thermo-PETase display comparable overall area of FELs, they are populated at different (meta)stable states. Apart from the shared states 1 and 5, where PETase exhibits the cleft-open, binding-competent structure, WT- and Thermo-PETase were also found to be populated at states 6 and 7, respectively. Detailed analysis reveals that WT-PETase maintains structural similarity at the catalytic site between states 1 and 6, indicating that WT-PETase in state 6 may not be advantageous for PET binding. On the other hand, Thermo-PETase in state 7 features an open-binding cleft with the elongated pairs of Ser160–His237 and Asp206– His237, thus rendering it a binding-competent state for promoting PET binding. In addition, it is evident that ThermoPETase can explore more conformational space at large distance values of Asp206–His237. It is worth noting that the Asp206–His237 bridge serves as a critical structural scaffold for the substrate accommodation and directly interacts with PET molecules (Figure 1I and Figure S1). In this respect, the extended and flexible Asp206–His237 pair possessed by Thermo-PETase may assist PET binding through excessive structural fluctuations.

### 2.3 Frustrations in native structures of PETase

A naturally evolved, well-folded protein adheres to the “principle of minimal frustration” [48], which typically results in a funneled energy landscape. However, conflicting inter-residue interactions, known as frustrated contacts, are frequently observed in the native structures of proteins. Although these interactions generally weaken the stability of native structures, they can promote specific structural dynamics that may be related to functional purposes [49]. To assess the presence of frustrated contacts in WT- and Thermo-PETase, we quantified frustrations in the native structures of these two PETases using the method introduced by Ferreiro et al. [50]. In brief, this method compares the energetic contribution to the additional stabilization provided to a pair of residues in the native structure with the statistical distribution of energies that would result from placing different residues in the same position. A native pair is deemed a highly frustrated contact if it induces significant destabilization compared to other possibilities, and vice versa for a minimally frustrated contact. The level of frustration is measured by a quantity, named the frustration index (*FruInd*), where a high (low) value of *FruInd* indicates a low (high) degree of frustration for the contact.

We observed wide distributions of *FruInd* for both WT- and Thermo-PETase, with highly frustrated contacts being only weakly populated (Figure 4A, 4B, 4C and Table S1). Additionally, the distributions of *FruInd* for WT- and Thermo-PETase largely overlap, suggesting that the three mutations in ThermoPETase may not lead to significant changes in the frustrations of the native structures. These results imply that both PETase and its variant have largely eliminated the highly frustrated interactions in their native states, resulting in stable structures as robust scaffolds for function. Focusing on the PET binding site, we observed that the distributions of *FruInd* for WT- and Thermo-PETase slightly shift to the left, indicating an accumulation of highly frustrated contacts at the binding site (Figure 4D). This suggests that the binding site of PETase is fragile and prone to deformation [51]. Furthermore, notable differences in frustration were observed between WT- and ThermoPETase (Figure 4E and 4F). Detailed comparisons at the binding site reveal that Thermo-PETase exhibits lower values in the *FruInd* distribution and possesses more highly frustrated contacts than WT-PETase. These findings together suggest that WT- and Thermo-PETase may share globally similar structural dynamics, with discrepancies primarily observed locally at the binding site, where Thermo-PETase displays more significant structural flexibility than WT-PETase.

**Figure 4:**
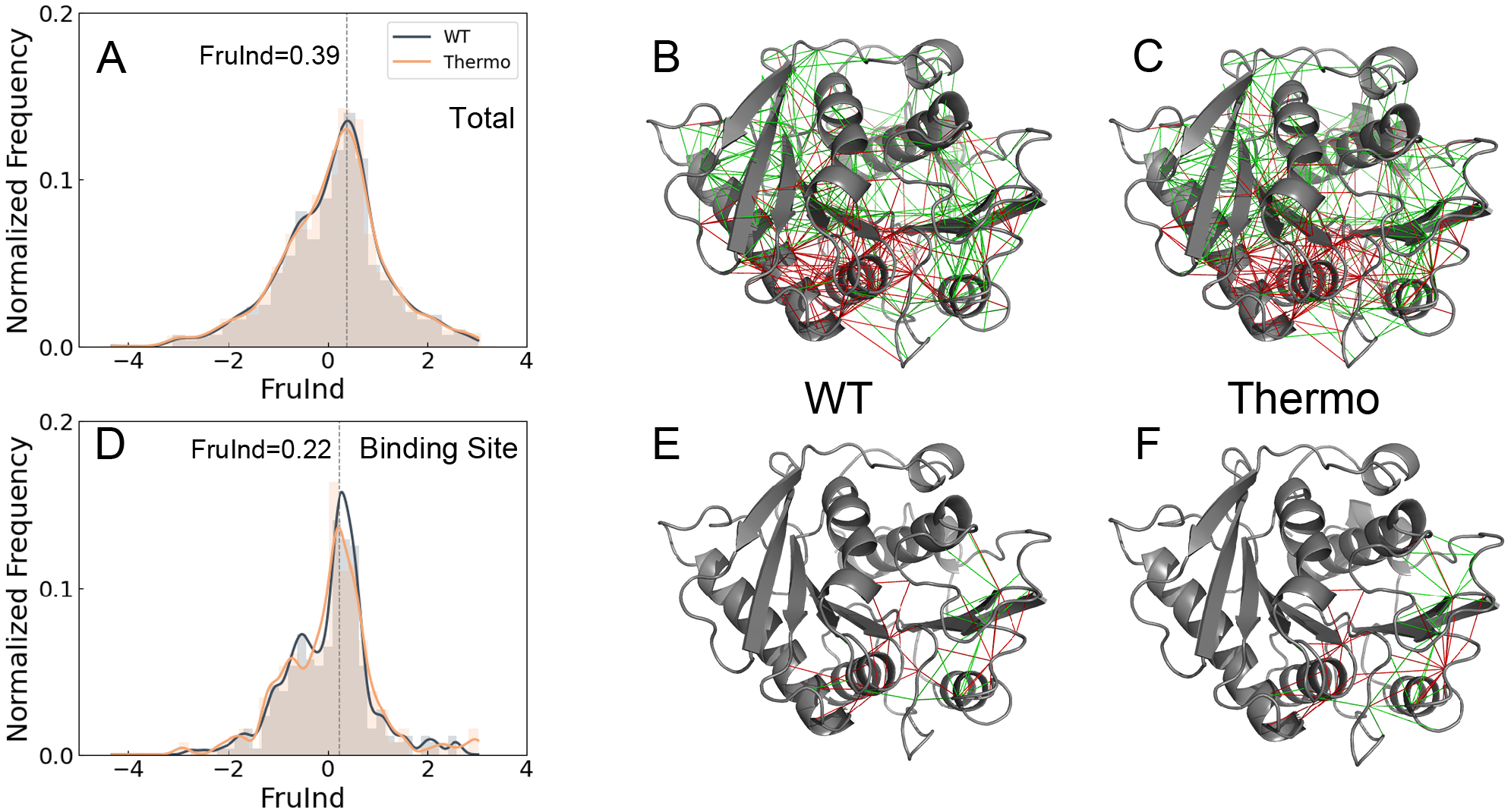
Local frustrations in WT- and Thermo-PETase. (A) Distribution of the frustration index (*FruInd*) for contacts involving all residues, with a dashed grey line indicating the most probable value. (B) Distribution of *FruInd* specifically for contacts involving residues within the binding site, with a dashed grey line indicating the most probable value. Visualization of total frustration on the structures of (C) WT- and (D) Thermo-PETase, respectively. Visualization of frustration at the binding site on the structures of (E) WT- and (F) Thermo-PETase, respectively. In (C), (D), (E) and (F), green lines represent the minimally frustrated contacts, while red lines represent the highly frustrated contacts. Neutral contacts are not shown.

### 2.4 Dynamics of PETase upon PET binding

Structural flexibility of enzymes at catalytic sites can be advantageous for substrate recruitment and product release [52, 53], however excessive structural dynamics at catalytic sites may induce large catalytic distances, thereby impeding chemical reactions. In order to see how PET binding impacts the structural dynamics of PETase and subsequent catalysis, we performed molecular docking of a PET tetramer (4PET) with PETase (Figure S1). The complex structure of PET:PETase obtained through docking is consistent with those from the previous studies [19, 54], and subsequently was used as the initial structure for the MD simulations at ambient temperatures (298K and 308K).

It has been proposed that the nucleophilic attack carried out by Ser160 to the carbonyl group of the benzene ring of the substrate PET molecules is essential for initializing the hydrolysis of PET polymer by PETase [37, 55, 56]. In this regard, we calculated the distance (*d*_*OC*_) between the oxygen *O*_*γ*_ of the catalytic Ser160 and the carbonyl carbon *C* of the substrate during MD simulations (Figure S6), with the fact that a chemical reaction only occurs when the involving atoms are spatially close. At 298K, both WT- and Thermo-PETase keep the distance *d*_*OC*_ at relatively low values, optimal for the subsequent catalytic reaction. When the temperature increases to 308K, the distance *d*_*OC*_ during the simulations largely remained at small values. We note that the large fluctuations in *d*_*OC*_ were observed occasionally (*d*_*OC*_ *>* 6 *Å*) at both temperatures, leading to substrate drifting away from the binding site. Due to the inherent flexible characteristics of the polymer, the enzyme-substrate bound state was observed to be flexible in our simulations, consistent with previous studies [54, 57].

In order to assess the structural dynamics of PETase upon substrate binding, we extracted the trajectories when 4PET is bound with PETase and calculated the RMSF of PETase (Figure 5A, 5B and S7). Interestingly, we found that the RMSF profiles of WT- and Thermo-PETase in the PET-bound binary complex are very similar. This is different from PETase at the apo state, where arresting discrepancies were found upon mutations (Figure 1). Detailed comparisons of RMSF profiles between WT- and Thermo-PETase indicate that Thermo-PETase has overall smaller RMSF values than WT-PETase, in particular at the catalytic residues of Asp206 and His237. Further structural analysis reveals that the catalytic triad of PETase in the bound complex is very similar for bothe WT- and ThermoPETase (Figure S8), resembling that in the native structure of PETase in its apo state. Our findings indicate that WT- and Thermo-PETase share similar structures and dynamics when they are bound with substrate.

We observed extensive quenching of structural dynamics in PETase bound with 4PET (Figure 5C and 5D). In detail, the most significant dynamics quenching effects for WT-PETase occurred within the W-loop. The side chain of Trp185 has been found to adopt three conformations in the native structure of WT-PETase [37, 38]. Quenching the structural dynamics of Trp185 and adapting its structure towards the one in the PET-binding state are crucial steps in realizing the catalysis of PETase, leading to the major difference between mesophilic and thermophilic PET hydrolases. On the other hand, 4PET significantly stabilizes the catalytic residues

**Figure 5:**
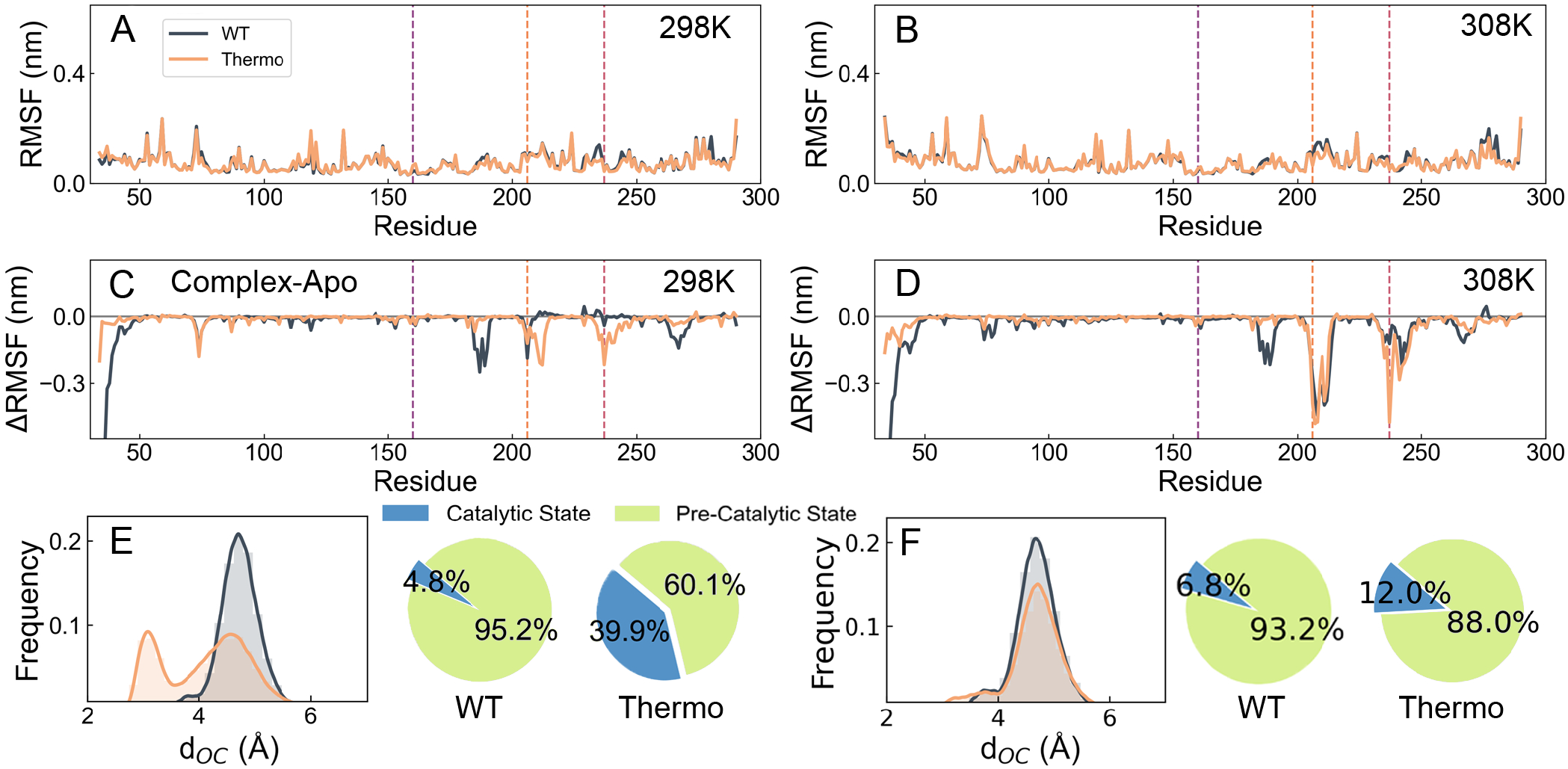
Structural dynamics of PETase upon substrate binding. RMSF of WT- and Thermo-PETase binding with substrate at (A) 298K and (B) 308K. RMSF differences of PETase between the apo state and the binary substrate-complex state for WT- and Thermo-PETase at (C) 298K and (D) 308K. (E) Normalized frequency distribution of spatial distance *d*_*OC*_ between the oxygen *O*_*γ*_ and the carbonyl carbon *C* of the substrate during MD simulations at (E) 298K and (F) 308K. The catalytic state and pre-catalytic state are defined by a threshold of *d*_*OC*_ of 4.0 *Å*.

Asp206 and His237 in Thermo-PETase, thus the dynamics of catalytic triad observed in apo state has been largely quenched. As the stable catalytic sites and spatially closed catalytic distances are prerequisite for catalysis, 4PET binding stabilizes structures of Thermo-PETase more than those of WT-PETase, possibly due to the fact that the interactions formed between substrate and enzyme are stronger and more widely distributed in Thermo-PETase than WT-PETase (Figure S9).

By analyzing the *d*_*OC*_ distribution, we found that ThermoPETase has a closer catalytic distance when bound to 4PET, leading to a higher population at the catalytic state, than WT-PETase (Figure 5E and 5F). This implies that Thermo-PETase is easier to trigger the subsequent catalytic reaction than WT-PETase, potentially contributing to the enhanced efficiency of catalysis. As the temperature increased from 298K to 308K, we found a decreasing population of the catalytic state for Thermo-PETase while WT-PETase maintained similar distributions of the catalytic and pre-catalytic states. Our results suggest that the catalytic sites of Thermo-PETase, compared to those of WT-PETase, are more sensitive to external stimuli, such as substrate binding and temperature changes.

### 3 Discussion and conclusions

In this work, we used MD simulations combined with local frustration analysis to investigate how the three mutations (S121E/D186H/R280A) contributed to the enhanced catalysis efficiency of Thermo-PETase. Although the initial design objective for Thermo-PETase was to increase the thermostability of the enzyme by introducing additional stabilizing interactions [21], we observed significant enhancements of structural flexibility at the PET binding sites. The open and dynamic nature of the active-site cleft has been recognized as the critical factor enabling PETase to function at ambient temperatures, providing a distinct catalytic advantage over thermophilic hydrolases [38, 40]. Our analysis of local frustration shows consistent results with MD simulations that introduction of the three mutations in Thermo-PETase increases the degree of frustration at the local PET binding sites, thus making Thermo-PETase be prone to deformation in favor of accommodating the substrate during binding. Noteworthy, our results on apo form of PETase reveal the dynamic picture of PETase when recruiting PET substrate, reminiscent of recently proposed conformational selection for PETase:PET complex initiation [46]. Interestingly, we found that substrate binding significantly reduces the structural dynamics of ThermoPETase induced by the mutations, resulting in a stable catalytic triad, similar to that of WT-PETase. Furthermore, we observed increases in substrate-enzyme inter-chain contacts in ThermoPETase, leading to more populated catalytic states for chemical reactions compared to WT-PETase. Collectively, we propose that the effects of mutations in Thermo-PETase on improving the PET degradation performance may be multifaceted: (1) enhancing the overall stability of the enzyme for increasing its tolerance to the environmental changes; (2) increasing the local structural flexibility at the binding site, which is advantageous for substrate recruitment and product release; (3) quenching the structural dynamics of enzyme at catalytic sites upon substrate binding, a perquisite for chemical reactions.

Substrate-binding residues are important for enzymatic activity. Previous studies identified a unique conformationally dynamical equilibrium of the Trp185 side chain wobbling between conformers A, B and C in PETase [37, 38]. This has made PETase distinguishable from other homologous enzymes, such as thermophilic PET hydrolases, where the equivalent conserved Trp adopts conformer C (Figure S10). Structural analysis indicates that substrate can only bind to PETase with Trp185 in conformer B by forming stacked interactions with one of the rings in PET molecules, while other conformers of Trp185 would clash with the substrate, thus hindering the binding [37]. Recent experiments and simulations have underlined the importance of the unique amino acids Ser214 and ILe218 of WT-PETase in triggering the high mobility of Trp185, thereby proposing a general mechanism for improving the catalytic activity of thermophilic PET hydrolases through the permission of the “wobbling” dynamics of Trp185 [45, 58]. Interestingly, our simulation results reveal that the conformational dynamics of Trp185 in Thermo-PETase have vanished, associated with a stabilized W-loop, compared to WT-PETase. Detailed structural analysis shows that Trp185 in Thermo-PETase is already in conformer B (Figure S10), thus facilitating the subsequent substrate binding. It is worth noting that great efforts have been made recently on mutating residues in the W-loop, aiming at fixing Trp185 in conformer B while simultaneously enhancing the overall stability [32, 26, 33]. Consequently, the mutations in Thermo-PETase have effectively stabilized both the local structure at Trp185 towards the substrate-binding conformation and the global structure by introducing a hydrogen bond (S121E/D186H) at the Wloop, which is one of the most flexible regions in WT-PETase [21, 59].

Conventionally, the practical design of WT-PETase aims to improve the thermostability by reducing the structural flexibility [21, 27, 60, 32, 26, 25, 29, 33, 24, 30], as stability and flexibility are usually considered to be coupled in proteins. For instance, the residues within the highly flexible W-loop of WT-PETase have been targeted as the hot spots for engineering. Through the substitutions of Asp186 with polar or non-polar residues in the W-loop [21, 32, 33], most PETase variants exhibited increased melting temperatures along with enhanced catalytic activity, compared to WT-PETase. From the energy landscape perspective, the stability of a protein can be associated with the degree of funnelness of the energy landscape [61, 62]. In this regard, mutations aiming at introducing stabilized interactions at the native state, which corresponds to the bottom of the funnel, will deepen the energy landscape, thus leading to increased thermostability (Figure 6). Interestingly, our simulations reveal that the improvements in the catalytic activity of Thermo-PETase can be partially attributed to the enhanced structural flexibility of the enzyme at its native state. In other words, in addition to increasing the depth of the funnel, mutations in Thermo-PETase have unprecedented effects on modulating the shape of the energy landscape at the bottom of the funnel. At low temperature, ThermoPETase can explore a wider range of conformational space than WT-PETase. Mutations enable Thermo-PETase to switch between the native structure resembling WT-PETase and the PET binding-competent structure with the open and flexible active cleft, which promotes substrate binding. This binding-competent structure can be highly populated for WT-PETase only at elevated temperatures, where Thermo-PETase becomes more flexible in rendering multiple binding-competent structures. However, WT-PETase is heat-labile and prone to denaturation, slowing down its catalytic activity at high temperatures. Upon PET binding, the extensive structural dynamics of Thermo-PETase are largely quenched by PET, leading to the stable catalytically competent native structure, same as WT-PETase. Furthermore, we found that the complex formed by WT-PETase and PET switch between catalytic and precatalytic states, supported by the NMR experiments, where PET in the binding cleft of PETase was found to be highly dynamic [39, 63]. Compared to WT-PETase, Thermo-PETase, which forms more extensive contacts with PET and is dynamically quenched, exhibits closer catalytic distances for nucleophilic attack, thus leading to higher activity for PET hydrolysis. In conclusion, mutations in Thermo-PETase have decoupled the interplay between global stability and local flexibility of the enzyme, thus improving the catalytic performance of Thermo-PETase over WT-PEase across a wide range of temperatures [21].

**Figure 6:**
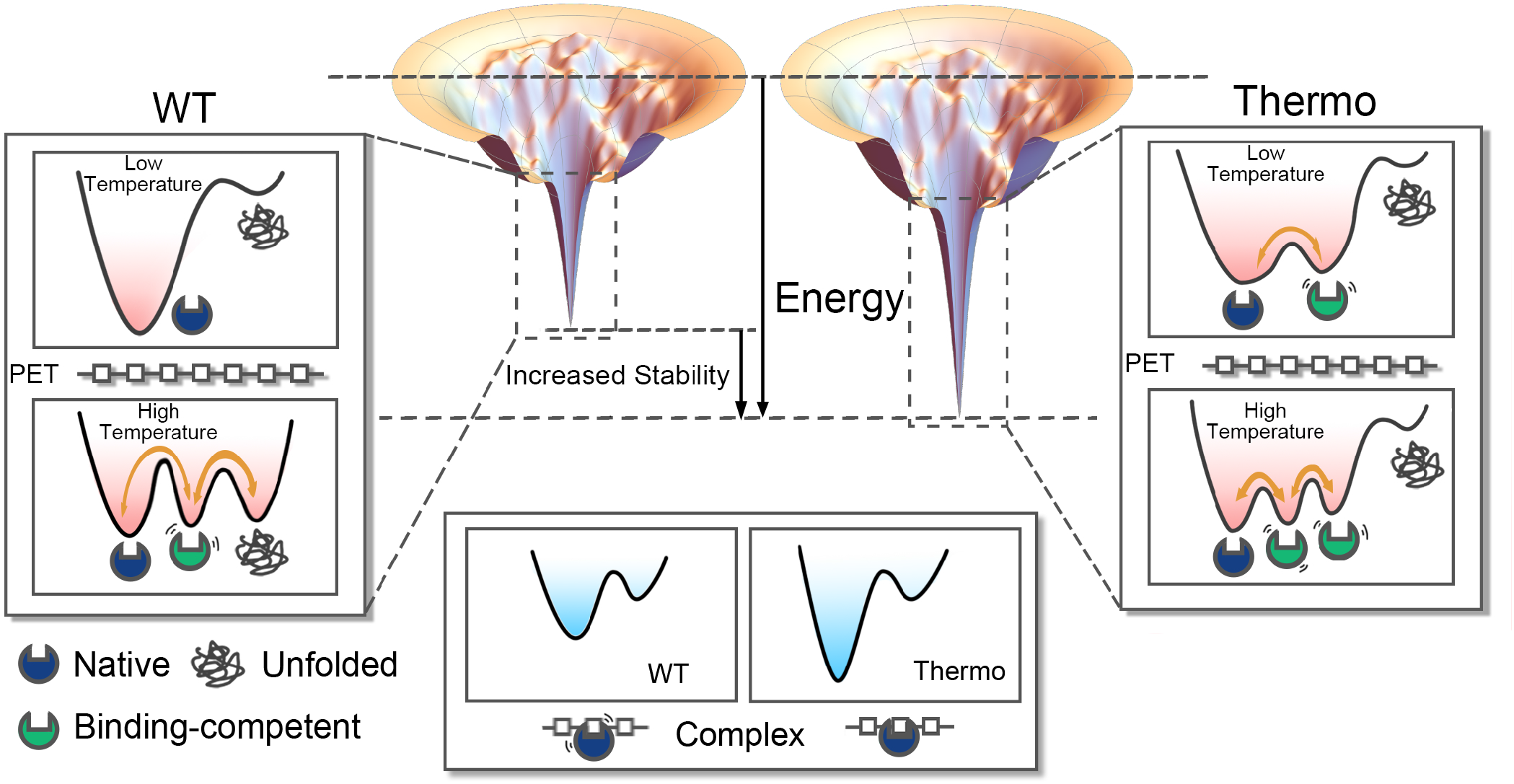
Energy landscapes and FELs of WT- and Thermo-PETase. The funneled energy landscapes of WT- and Thermo-PETase show arresting differences in the depth of funnels (*upper center*), suggesting increased stability for Thermo-PETase upon the triple mutations. With close examination at the bottom of the funnel, WT- and Thermo-PETase exhibit distinct free energy profiles (*left* and *right*), each showcasing unique temperature responses. Orange arrows denote the transitions between states near the native state, with arrow thickness representing the ease of these transitions. Upon PET binding (*lower center*), these two PETases have the same catalytically competent native structure with PET being more loosely bound with WT-PETase than Thermo-PETase.

In summary, our results reveal that the stability and flexibility of enzymes are not necessarily correlated, in contradiction to the convention that high (low) stability usually leads to a rigid (flexible) protein structure [64]. It is worth noting that a recent work has observed a decoupling interplay between the stability and flexibility in a computationally designed hydrolase Turbo-PETase, which has demonstrated superior performance compared to other PET hydrolases to date [30]. The *N* high PET depolymerization efficiency of Turbo-PETase was ^*N*^ attributed to the mutations that have resulted in both increases in thermostability at the global scale and structural flexibility at the PET-binding site, exhibiting elevated melting temperature and simultaneously enabling the promiscuous attack to different PET surface structures [30]. This significant achievement made by Turbo-PETase has paved a promising way for future PETase design, focusing on enhancing both thermostability and flexibility. Although the full mechanistic understanding of achieving an optimal balance between the stability and flexibility in PETase for highly efficient PET degradation remains elusive, our study offers new insights into the rational design strategy to improve the catalytic activity and thermostability of PETase, where both the structure and dynamics of the enzyme should be simultaneously considered.

## 4 Materials and Methods

### 4.1 Molecular dynamics simulations

We performed all-atom MD simulations using Gromacs2023.2 software [65]. MD simulations were conducted with the Amber ff14SB force field in conjunction with the TIP3P water model [66]. Simulations were initialized from the respective native structures for WT- and Therm-PETase (PDB: 5XJH [19] and 6IJ6 [21]). The native structures of WT- and Thermo-PETase were individually placed in a cubic box with margins of 10 *Å*. The protein systems were solvated with the TIP3P water model and the salt concentration was set to be 100 *nM* to mimick the *in vivo* environment. A short energy minimization step in each simulation system was done through the steepest descent algorithm. Systems were then equilibrated at the NVT phase using the v-rescale thermostat. Subsequently, the NPT simulations were conducted with the Parrinello-Rahman barostat at a relaxation time of *τ*_*p*_ = 4.0 *ps* [67]. The LINCS algorithm was used to constrain all hydrogen bonds [68], resulting in a time step of 2 *f s*. The particle-mesh Ewald (PME) approach for computing the long-range electrostatic interactions was used [69] and all the non-bonded interactions were cut off at 10 *Å*. Simulations were run for 10 *μ*s at the ambient temperatures (298K and 308K) and 1 *μ*s at an elevated temperature of 450K. For each PETase system, two trajectories were performed at every simulation temperature.

### 4.2 Trajectory analysis

We used the built-in modules of Gromacs to calculate RMSD, RMSF, residue distances and conduct PCA, while MDAnalysis was employed for calculating the contact map [70]. All the analyses were performed after removal of the translation and rotation of the protein in the simulation system during the trajectories. RMSF value for each residue was calculated by averaging RMSF of atoms belonging to that residue. For contact probability calculations, contacts between any heavy atoms of one residue pair were included with the cut-off distance set at 5Å. The results were normalized to obtain the residue-level contact probability. For PCA, the covariance value *C*_*i j*_ between residue *i* and *j* was firstly calculated by: 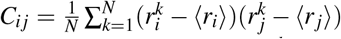, where *N* is the total number of simulation frames, 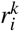 is the coordinate of *C*_*α*_ atom of residue *i* in frame *k*. Then, the eigenvectors (PCs) and eigenvalues were calculated by diagonalizing the covariance matrix. Frustration results were obtained by using the Frustratometer online server [71].

### 4.3 Molecular docking

We constructed the complex structure of PETase with a PET tetramer (4PET) by molecular docking. We used the crystal structure of WT-PETase as the receptor in docking (PDB: 5XJH [19]) and the 4PET ligand was built based on a PET dimer using Avogadro software [72, 73]. All hydrogen atoms were added to both the protein and ligand using AutoDockTools prior to performing docking [74]. Molecular docking was then carried out between the WT-PETase structure and 4PET by AutoDock Vina (version 1.1.2) [75]. During docking, the ligand was set to be flexible while the protein was kept rigid. To prevent the collapse of the PET chain, the three bonds within O–C–C–O connecting the second and third benzene rings and the central C–C bond connecting the first and second benzene rings of the PET ligand were not allowed to rotate. The docking model, which is consistent with those from previous studies [19, 54], was selected and further evaluation of the structure model was done by MD simulations. MD simulations of the PETase:4PET complex starting from the docking model were performed using Gromacs-2023.2 software, following the same protocols as applied in simulations of the apo state of PETase. PET ligand was parameterized with the general Amber force field (GAFF2) [76] and the topology files of 4PET for Gromacs simulations were prepared using ACPYPE software [77]. To model the complex structure of ThermoPETase with 4PET, we directly implemented three mutations on WT-PETase using PyMOL software [78]. For each PETase system, two trajectories with a length of 1 *μ*s were generated at both 298K and 308K.

## Data Availability

The necessary files for setting up Gromacs simulations and molecular docking, as well as the frustration analysis results are publicly available at https://osf.io/9yzjf/. Additional data can be found in Supporting Information.

## Acknowledgement

The project was supported by National Key R&D Program of China (No. 2023YFC3905000) and Guangdong Scientific Research Platform and Projects for Higher-educational Institutions from Department of Education of Guangdong Province (Grant No. 2023KTSCX169). X.C. was also partly supported by the Municipal Key Laboratory Construction program of the Guangzhou Municipal Science and Technology Project (Grant No. 2023A03J0003).

## References

[1] Rucha V. Moharir and Sunil Kumar. Challenges associated with plastic waste disposal and allied microbial routes for its effective degradation: A comprehensive review. Journal of Cleaner Production, 208:65–76, January 2019.

[2] P. O. Awoyera and A. Adesina. Plastic wastes to construction products: Status, limitations and future perspective. Case Studies in Construction Materials, 12:e00330, June 2020.

[3] Sameh Samir Ali, Tamer Elsamahy, Eleni Koutra, Michael Kornaros, Mostafa El-Sheekh, Esraa A. Abdelkarim, Daochen Zhux, and Jianzhong Sun. Degradation of conventional plastic wastes in the environment: A review on current status of knowledge and future perspectives of disposal. Science of The Total Environment, 771:144719, June 2021.

[4] Jenna R Jambeck, Roland Geyer, Chris Wilcox, Theodore R Siegler, Miriam Perryman, Anthony Andrady, Ramani Narayan, and Kara Lavender Law. Plastic waste inputs from land into the ocean. Science, 347(6223):768–771, 2015.

[5] Roland Geyer, Jenna R Jambeck, and Kara Lavender Law. Production, use, and fate of all plastics ever made. Science advances, 3(7):e1700782, 2017.

[6] Vijaykumar Sinha, Mayank R Patel, and Jigar V Patel. Pet waste management by chemical recycling: a review. Journal of Polymers and the Environment, 18(1):8–25, 2010.

[7] Kara Lavender Law, Natalie Starr, Theodore R. Siegler, Jenna R. Jambeck, Nicholas J. Mallos, and George H. Leonard. The united states’ contribution of plastic waste to land and ocean. Science Advances, 6(44):eabd0288, October 2020.

[8] Ren Wei and Wolfgang Zimmermann. Biocatalysis as a green route for recycling the recalcitrant plastic polyethylene terephthalate. Microbial biotechnology, 10(6):1302, 2017.

[9] Athena Papadopoulou, Katrin Hecht, and Rebecca Buller. Enzymatic pet degradation. Chimia, 73(9):743–743, 2019.

[10] Ikuo Taniguchi, Shosuke Yoshida, Kazumi Hiraga, Kenji Miyamoto, Yoshiharu Kimura, and Kohei Oda. Biodegradation of pet: current status and application aspects. Acs Catalysis, 9(5):4089–4105, 2019.

[11] Fusako Kawai, Takeshi Kawabata, and Masayuki Oda. Current knowledge on enzymatic pet degradation and its possible application to waste stream management and other fields. Applied microbiology and biotechnology, 103:4253–4268, 2019.

[12] Yunyi Wei, Lora Swenson, Clementina Castro, Urszula Derewenda, Wladek Minor, Hiroyuki Arai, Junken Aoki, Keizo Inoue, Luis Servin-Gonzalez, and Zygmunt S Derewenda. Structure of a microbial homologue of mammalian platelet-activating factor acetylhydrolases: Streptomyces exfoliatus lipase at 1.9 å resolution. Structure, 6(4):511–519, 1998.

[13] Rolf-Joachim Müller, Hedwig Schrader, Jörn Profe, Karolin Dresler, and Wolf-Dieter Deckwer. Enzymatic degradation of poly(ethylene terephthalate): Rapid hydrolyse using a hydrolase from t. fusca. Macromolecular Rapid Communications, 26(17):1400–1405, 2005.

[14] Åsa M Ronkvist, Wenchun Xie, Wenhua Lu, and Richard A Gross. Cutinase-catalyzed hydrolysis of poly (ethylene terephthalate). Macromolecules, 42(14):5128– 5138, 2009.

[15] Sintawee Sulaiman, Saya Yamato, Eiko Kanaya, Joong-Jae Kim, Yuichi Koga, Kazufumi Takano, and Shigenori Kanaya. Isolation of a novel cutinase homolog with polyethylene terephthalate-degrading activity from leaf-branch compost by using a metagenomic approach. Applied and Environmental Microbiology, 78(5):1556– 1562, 2012.

[16] Ren Wei, Daniel Breite, Chen Song, Daniel Gräsing, Tina Ploss, Patrick Hille, Ruth Schwerdtfeger, Jörg Matysik, Agnes Schulze, and Wolfgang Zimmermann. Biocatalytic degradation efficiency of postconsumer polyethylene terephthalate packaging determined by their polymer microstructures. Advanced Science, 6(14):1900491, 2019.

[17] V Tournier, CM Topham, A Gilles, B David, C Folgoas, E Moya-Leclair, E Kamionka, M-L Desrousseaux, H Texier, S Gavalda, et al. An engineered pet depolymerase to break down and recycle plastic bottles. Nature, 580(7802):216–219, 2020.

[18] Shosuke Yoshida, Kazumi Hiraga, Toshihiko Takehana, Ikuo Taniguchi, Hironao Yamaji, Yasuhito Maeda, Kiyotsuna Toyohara, Kenji Miyamoto, Yoshiharu Kimura, and Kohei Oda. A bacterium that degrades and assimilates poly(ethylene terephthalate). Science, 351(6278):1196– 1199, March 2016.

[19] Seongjoon Joo, In Jin Cho, Hogyun Seo, Hyeoncheol Francis Son, Hye-Young Sagong, Tae Joo Shin, So Young Choi, Sang Yup Lee, and Kyung-Jin Kim. Structural insight into molecular mechanism of poly(ethylene terephthalate) degradation. Nature Communications, 9(1):382, December 2018.

[20] Erika Erickson, Thomas J Shakespeare, Felicia Bratti, Bonnie L Buss, Rosie Graham, McKenzie A Hawkins, Gerhard König, William E Michener, Joel Miscall, Kelsey J Ramirez, et al. Comparative performance of petase as a function of reaction conditions, substrate properties, and product accumulation. ChemSusChem, 15(1):e202101932, 2022.

[21] Hyeoncheol Francis Son, In Jin Cho, Seongjoon Joo, Hogyun Seo, Hye-Young Sagong, So Young Choi, Sang Yup Lee, and Kyung-Jin Kim. Rational protein engineering of thermo-stable petase from ideonella sakaiensis for highly efficient pet degradation. ACS Catalysis, 9(4):3519–3526, April 2019.

[22] Xiangxi Meng, Lixin Yang, Hanqing Liu, Qingbin Li, Guoshun Xu, Yan Zhang, Feifei Guan, Yuhong Zhang, Wei Zhang, Ningfeng Wu, et al. Protein engineering of stable ispetase for pet plastic degradation by premuse. International Journal of Biological Macromolecules, 180:667–676, 2021.

[23] Yidi Liu, Zhanzhi Liu, Zhiyong Guo, Tingting Yan, Changxu Jin, and Jing Wu. Enhancement of the degradation capacity of ispetase for pet plastic degradation by protein engineering. Science of The Total Environment, 834:154947, August 2022.

[24] Yinglu Cui, Yanchun Chen, Xinyue Liu, Saijun Dong, Yu’e Tian, Yuxin Qiao, Ruchira Mitra, Jing Han, Chunli Li, Xu Han, Weidong Liu, Quan Chen, Wangqing Wei, Xin Wang, Wenbin Du, Shuangyan Tang, Hua Xiang, Haiyan Liu, Yong Liang, Kendall N. Houk, and Bian Wu. Computational redesign of a petase for plastic biodegradation under ambient condition by the grape strategy. ACS Catalysis, 11(3):1340–1350, February 2021.

[25] Hongyuan Lu, Daniel J. Diaz, Natalie J. Czarnecki, Congzhi Zhu, Wantae Kim, Raghav Shroff, Daniel J. Acosta, Bradley R. Alexander, Hannah O. Cole, Yan Zhang, Nathaniel A. Lynd, Andrew D. Ellington, and Hal S. Alper. Machine learning-aided engineering of hydrolases for pet depolymerization. Nature, 604(7907):662–667, April 2022.

[26] Elizabeth L Bell, Ross Smithson, Siobhan Kilbride, Jake Foster, Florence J Hardy, Saranarayanan Ramachandran, Aleksander A Tedstone, Sarah J Haigh, Arthur A Garforth, Philip JR Day, et al. Directed evolution of an efficient and thermostable pet depolymerase. Nature Catalysis, 5(8):673–681, 2022.

[27] Stefan Brott, Lara Pfaff, Josephine Schuricht, Jan-Niklas Schwarz, Dominique Böttcher, Christoffel PS Badenhorst, Ren Wei, and Uwe T Bornscheuer. Engineering and evaluation of thermostable ispetase variants for pet degradation. Engineering in life sciences, 22(3-4):192– 203, 2022.

[28] Seul Hoo Lee, Hogyun Seo, Hwaseok Hong, Jiyoung Park, Dongwoo Ki, Mijeong Kim, Hyung-Joon Kim, and Kyung-Jin Kim. Three-directional engineering of ispetase with enhanced protein yield, activity, and durability. Journal of Hazardous Materials, 459:132297, 2023.

[29] Lixia Shi, Pi Liu, Zijian Tan, Wei Zhao, Junfei Gao, Qun Gu, Hongwu Ma, Haifeng Liu, and Leilei Zhu. Complete depolymerization of pet wastes by an evolved pet hydrolase from directed evolution. Angewandte Chemie International Edition, 62(14):e202218390, 2023.

[30] Yinglu Cui, Yanchun Chen, Jinyuan Sun, Tong Zhu, Hua Pang, Chunli Li, Wen-Chao Geng, and Bian Wu. Computational redesign of a hydrolase for nearly complete pet depolymerization at industrially relevant high-solids loading. Nature Communications, 15(1):1417, February 2024.

[31] Yvonne Joho, Santana Royan, Alessandro Caputo, Sophia Newton, Thomas Peat, Janet Newman, Colin Jackson, and Albert Ardevol. Enhancing pet degrading enzymes: A combinatory approach. ChemBioChem, page e202400084, 2024.

[32] Qingdian Yin, Shengping You, Jiaxing Zhang, Wei Qi, and Rongxin Su. Enhancement of the polyethylene terephthalate and mono-(2-hydroxyethyl) terephthalate degradation activity of ideonella sakaiensis petase by an electrostatic interaction-based strategy. Bioresource Technology, 364:128026, 2022.

[33] Qingdian Yin, Jiaxing Zhang, Sen Ma, Tao Gu, Mengfan Wang, Shengping You, Sheng Ye, Rongxin Su, Yaxin Wang, and Wei Qi. Efficient polyethylene terephthalate biodegradation by an engineered ideonella sakaiensis petase with a fixed substrate-binding w156 residue. Green Chemistry, 26:2560–2570, 2024.

[34] Ren Wei, Gerlis von Haugwitz, Lara Pfaff, Jan Mican, Christoffel PS Badenhorst, Weidong Liu, Gert Weber, Harry P Austin, David Bednar, Jiri Damborsky, et al. Mechanism-based design of efficient pet hydrolases. ACS catalysis, 12(6):3382–3396, 2022.

[35] Lita Amalia, Chia-Yu Chang, Steven SS Wang, Yi-Chun Yeh, and Shen-Long Tsai. Recent advances in the biological depolymerization and upcycling of polyethylene terephthalate. Current Opinion in Biotechnology, 85:103053, 2024.

[36] Lixia Shi and Leilei Zhu. Recent advances and challenges in enzymatic depolymerization and recycling of pet wastes. ChemBioChem, 25(2):e202300578, 2024.

[37] Xu Han, Weidong Liu, Jian-Wen Huang, Jiantao Ma, Yingying Zheng, Tzu-Ping Ko, Limin Xu, Ya-Shan Cheng, Chun-Chi Chen, and Rey-Ting Guo. Structural insight into catalytic mechanism of pet hydrolase. Nature communications, 8(1):2106, 2017.

[38] Harry P Austin, Mark D Allen, Bryon S Donohoe, Nicholas A Rorrer, Fiona L Kearns, Rodrigo L Silveira, Benjamin C Pollard, Graham Dominick, Ramona Duman, Kamel El Omari, et al. Characterization and engineering of a plastic-degrading aromatic polyesterase. Proceedings of the National Academy of Sciences, 115(19):E4350–E4357, 2018.

[39] Ren Wei, Chen Song, Daniel Gräsing, Tobias Schneider, Pavlo Bielytskyi, Dominique Böttcher, Jörg Matysik, Uwe T Bornscheuer, and Wolfgang Zimmermann. Conformational fitting of a flexible oligomeric substrate does not explain the enzymatic pet degradation. Nature communications, 10(1):5581, 2019.

[40] Tobias Fecker, Pablo Galaz-Davison, Felipe Engelberger, Yoshie Narui, Marcos Sotomayor, Loreto P. Parra, and César A. Ramírez-Sarmiento. Active site flexibility as a hallmark for efficient pet degradation by i. sakaiensis petase. Biophysical Journal, 114(6):1302–1312, March 2018.

[41] Carla Silva, Shi Da, Nádia Silva, Teresa Matamá, Rita Araújo, Madalena Martins, Sheng Chen, Jian Chen, Jing Wu, Margarida Casal, et al. Engineered thermobifida fusca cutinase with increased activity on polyester substrates. Biotechnology Journal, 6(10):1230–1239, 2011.

[42] Rita Araújo, Carla Silva, Alexandre ONeill, Nuno Micaelo, Georg Guebitz, Cláudio M Soares, Margarida Casal, and Artur Cavaco-Paulo. Tailoring cutinase activity towards polyethylene terephthalate and polyamide 6, 6 fibers. Journal of Biotechnology, 128(4):849–857, 2007.

[43] Yeyi Kan, Lihui He, Yunzi Luo, and Rui Bao. Ispetase is a novel biocatalyst for poly (ethylene terephthalate)(pet) hydrolysis. ChemBioChem, 22(10):1706–1716, 2021.

[44] Alessandro Berselli, Maria J Ramos, and Maria Cristina Menziani. Novel pet-degrading enzymes: Structure-function from a computational perspective. ChemBioChem, 22(12):2032–2050, 2021.

[45] Chun-Chi Chen, Xu Han, Xian Li, Pengcheng Jiang, D. Niu, Lixin Ma, Weidong Liu, Siyu Li, Yingying Qu, Hebing Hu, et al. General features to enhance enzymatic activity of poly (ethylene terephthalate) hydrolysis. Nature Catalysis, 4(5):425–430, 2021.

[46] Boyang Guo, Sudarsana Reddy Vanga, Ximena Lopez-Lorenzo, Patricia Saenz-Mendez, Sara Ronnblad Ericsson, Yuan Fang, Xinchen Ye, Karen Schriever, Eva Backstrom, Antonino Biundo, et al. Conformational selection in biocatalytic plastic degradation by petase. Acs Catalysis, 12(6):3397–3409, 2022.

[47] Diego U Ferreiro, Elizabeth A Komives, and Peter G Wolynes. Frustration, function and folding. Current Opinion in Structural Biology, 48:68–73, February 2018.

[48] Joseph D Bryngelson and Peter G Wolynes. Spin glasses and the statistical mechanics of protein folding. Proceedings of the National Academy of sciences, 84(21):7524– 7528, 1987.

[49] Diego U Ferreiro, Elizabeth A Komives, and Peter G Wolynes. Frustration in biomolecules. Quarterly reviews of biophysics, 47(4):285–363, 2014.

[50] Diego U. Ferreiro, Joseph A. Hegler, Elizabeth A. Komives, and Peter G. Wolynes. Localizing frustration in native proteins and protein assemblies. Proceedings of the National Academy of Sciences, 104(50):19819– 19824, December 2007.

[51] Mingchen Chen, Xun Chen, Nicholas P Schafer, Cecilia Clementi, Elizabeth A Komives, Diego U Ferreiro, and Peter G Wolynes. Surveying biomolecular frustration at atomic resolution. Nature communications, 11(1):5944, 2020.

[52] Gordon G Hammes. Multiple conformational changes in enzyme catalysis. Biochemistry, 41(26):8221–8228, 2002.

[53] Sharon Hammes-Schiffer and Stephen J Benkovic. Relating protein motion to catalysis. Annu. Rev. Biochem., 75:519–541, 2006.

[54] Clauber Henrique Souza da Costa, Alberto M Dos Santos, Cláudio Nahum Alves, Sérgio Martí, Vicent Moliner, Kaue Santana, and Jerônimo Lameira. Assessment of the petase conformational changes induced by poly (ethylene terephthalate) binding. Proteins: Structure, Function, and Bioinformatics, 89(10):1340–1352, 2021.

[55] Rafael García-Meseguer, Enrique Ortí, Iñaki Tuñón, J Javier Ruiz-Pernía, and Juan Aragó. Insights into the enhancement of the poly (ethylene terephthalate) degradation by fast-petase from computational modeling. Journal of the American Chemical Society, 145(35):19243–19255, 2023.

[56] Tucker Burgin, Benjamin C Pollard, Brandon C Knott, Heather B Mayes, Michael F Crowley, John E McGeehan, Gregg T Beckham, and H Lee Woodcock. The reaction mechanism of the ideonella sakaiensis petase enzyme. Communications Chemistry, 7(1):65, 2024.

[57] Linyu Chen, Fangfang Fan, Meiyuan Yang, Linquan Wang, Yushuo Bai, Shuai Qiu, Changjiang Lyu, and Jun Huang. Atomistic insight into the binding mode and self-regulation mechanism of is petase towards pet substrates with different polymerization degrees. Physical Chemistry Chemical Physics, 25(27):18332–18345, 2023.

[58] Alessandro Crnjar, Aransa Griñen, Shina C. L. Kamerlin, and César A. Ramírez-Sarmiento. Conformational selection of a tryptophan side chain drives the generalized increase in activity of pet hydrolases through a ser/ile double mutation. ACS Organic & Inorganic Au, 3(2):109– 119, April 2023.

[59] Zhi Qu, Lin Zhang, and Yan Sun. Molecular insights into the enhanced activity and/or thermostability of pet hydrolase by d186 mutations. Molecules, 29(6):1338, 2024.

[60] Hyeoncheol Francis Son, Seongjoon Joo, Hogyun Seo, Hye-Young Sagong, Seul Hoo Lee, Hwaseok Hong, and Kyung-Jin Kim. Structural bioinformatics-based protein engineering of thermo-stable petase from ideonella sakaiensis. Enzyme and Microbial Technology, 141:109656, 2020.

[61] Jin Wang, Ronaldo J Oliveira, Xiakun Chu, Paul C Whitford, Jorge Chahine, Wei Han, Erkang Wang, José N Onuchic, and Vitor BP Leite. Topography of funneled landscapes determines the thermodynamics and kinetics of protein folding. Proceedings of the National Academy of Sciences, 109(39):15763–15768, 2012.

[62] Xiakun Chu, Linfeng Gan, Erkang Wang, and Jin Wang. Quantifying the topography of the intrinsic energy landscape of flexible biomolecular recognition. Proceedings of the National Academy of Sciences, 110(26):E2342– E2351, 2013.

[63] Patricia Falkenstein, Ren Wei, Jörg Matysik, and Chen Song. Mechanistic investigation of enzymatic degradation of polyethylene terephthalate by nuclear magnetic resonance. In Methods in Enzymology, volume 648, pages 231–252. Elsevier, 2021.

[64] Tim J Kamerzell and C Russell Middaugh. The complex inter-relationships between protein flexibility and stability. Journal of pharmaceutical sciences, 97(9):3494– 3517, 2008.

[65] Mark James Abraham, Teemu Murtola, Roland Schulz, Szilárd Páll, Jeremy C. Smith, Berk Hess, and Erik Lindahl. Gromacs: High performance molecular simulations through multi-level parallelism from laptops to supercomputers. SoftwareX, 1–2:19–25, September 2015.

[66] James A. Maier, Carmenza Martinez, Koushik Kasavajhala, Lauren Wickstrom, Kevin E. Hauser, and Carlos Simmerling. ff14sb: Improving the accuracy of protein side chain and backbone parameters from ff99sb. Journal of Chemical Theory and Computation, 11(8):3696–3713, August 2015.

[67] M. Parrinello and A. Rahman. Polymorphic transitions in single crystals: A new molecular dynamics method. Journal of Applied Physics, 52(12):7182–7190, December 1981.

[68] Berk Hess, Henk Bekker, Herman J. C. Berendsen, and Johannes G. E. M. Fraaije. Lincs: A linear constraint solver for molecular simulations. Journal of Computational Chemistry, 18(12):1463–1472, 1997.

[69] Tom Darden, Darrin York, and Lee Pedersen. Particle mesh ewald: An n·log(n) method for ewald sums in large systems. The Journal of Chemical Physics, 98(12):10089–10092, June 1993.

[70] Richard J. Gowers, Max Linke, Jonathan Barnoud, Tyler J. E. Reddy, Manuel N. Melo, Sean L. Seyler, Jan Domański, David L. Dotson, Sébastien Buchoux, Ian M. Kenney, and Oliver Beckstein. Mdanalysis: A python package for the rapid analysis of molecular dynamics simulations. Proceedings of the 15th Python in Science Conference, pages 98–105, 2016.

[71] R. Gonzalo Parra, Nicholas P. Schafer, Leandro G. Radusky, Min-Yeh Tsai, A. Brenda Guzovsky, Peter G. Wolynes, and Diego U. Ferreiro. Protein frustratometer 2: A tool to localize energetic frustration in protein molecules, now with electrostatics. Nucleic Acids Research, 44(W1):W356–W360, July 2016.

[72] Carola Jerves, Rui PP Neves, Maria J Ramos, Saulo da Silva, and Pedro A Fernandes. Reaction mechanism of the pet degrading enzyme petase studied with dft/mm molecular dynamics simulations. ACS Catalysis, 11(18):11626–11638, 2021.

[73] Marcus D Hanwell, Donald E Curtis, David C Lonie, Tim Vandermeersch, Eva Zurek, and Geoffrey R Hutchison. Avogadro: an advanced semantic chemical editor, visualization, and analysis platform. Journal of cheminformatics, 4:1–17, 2012.

[74] Garrett M. Morris, Ruth Huey, William Lindstrom, Michel F. Sanner, Richard K. Belew, David S. Goodsell, and Arthur J. Olson. Autodock4 and autodocktools4: Automated docking with selective receptor flexibility. Journal of computational chemistry, 30(16):2785–2791, December 2009.

[75] Jerome Eberhardt, Diogo Santos-Martins, Andreas F. Tillack, and Stefano Forli. Autodock vina 1.2.0: New docking methods, expanded force field, and python bindings. Journal of Chemical Information and Modeling, 61(8):3891–3898, August 2021.

[76] Junmei Wang, Romain M. Wolf, James W. Caldwell, Peter A. Kollman, and David A. Case. Development and testing of a general amber force field. Journal of Computational Chemistry, 25(9):1157–1174, 2004.

[77] Alan W. Sousa da Silva and Wim F. Vranken. Acpype - antechamber python parser interface. BMC Research Notes, 5(1):367, July 2012.

[78] LLC Schrodinger. The pymol molecular graphics system. Version, 1:8, 2015.

