## Supplementary material for "Unraveling the Interplay between Stability and Flexibility in Design of Polyethylene Terephthalate (PET) Hydrolases": Supplymentary Information

[3] Division of Life Science
The Hong Kong University of Science and Technology
Clear Water Bay, Hong Kong SAR 999077, China

| | $N_{total}$ | $N_{BSite}$ | $N_{minimally}$ | Minimally% |
|---|---|---|---|---|
| WT | 2031 | 275 | 27 | 9.82 |
| Thermo | 2011 | 275 | 34 | 12.36 |
| | $N_{highly}$ | Highly% | | |
| WT | 35 | 12.73 | | |
| Thermo | 38 | 13.82 | | |

Table S1: Frustration analysis of WT- and Thermo-PETase. $N_{total}$ is the number of total contacts and $N_{BSite}$ is the number of contacts involving at least one residue within the PET binding site. $N_{minimally}$ and $N_{highly}$ are the number of minimally and highly frustrated contacts involving at least one residue within the PET binding site, respectively. Minimally% and Highly% are the percentages of minimally and highly frustrated contacts involving at least one residue within the PET binding site, respectively.

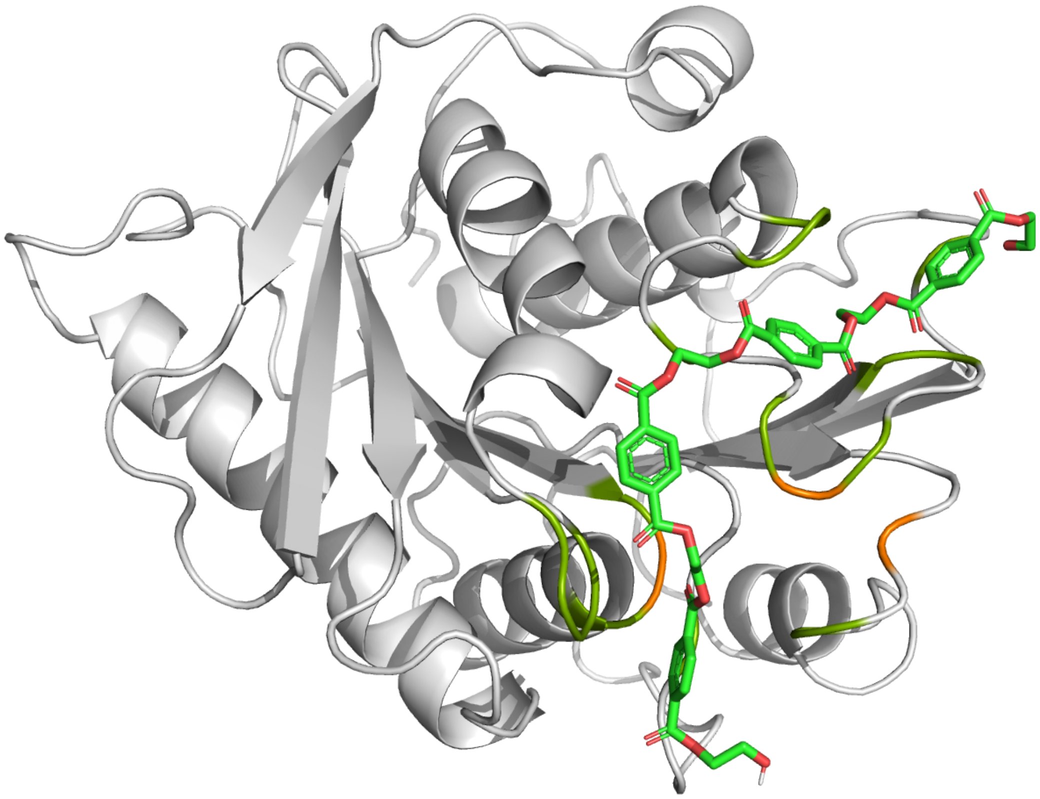

Figure S1: WT-PETase structure with a PET tetramer binding at the catalytic site. The molecular docking was set up using the AutoDock Tools [1]. The docking structure was obtained using the AutoDock Vina program [2]. The residues at the PET binding site are colored in dark green and the three catalytic residues (Ser160, Asp206 and His237) are colored in orange.

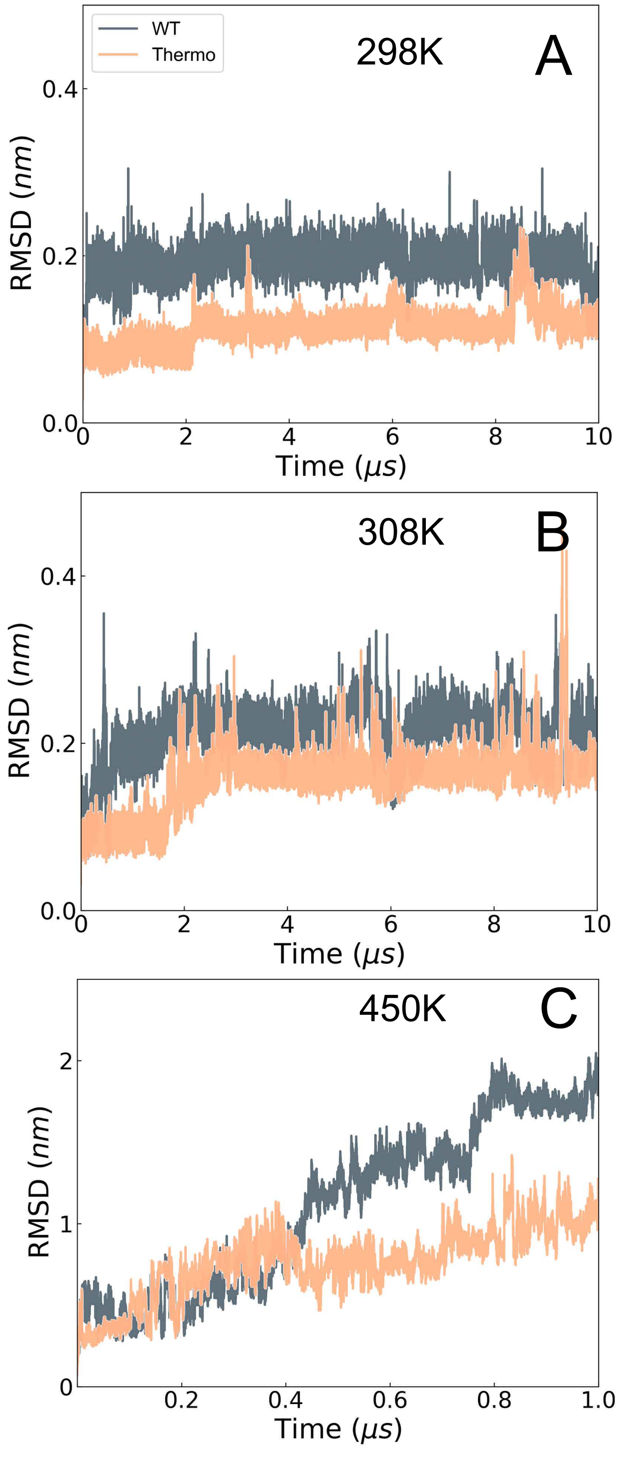


Figure S2: RMSD to the native PDB structures of the other simulation trajectories at each temperature of (A)298K, (B) 308K and (C) 450K, respectively.

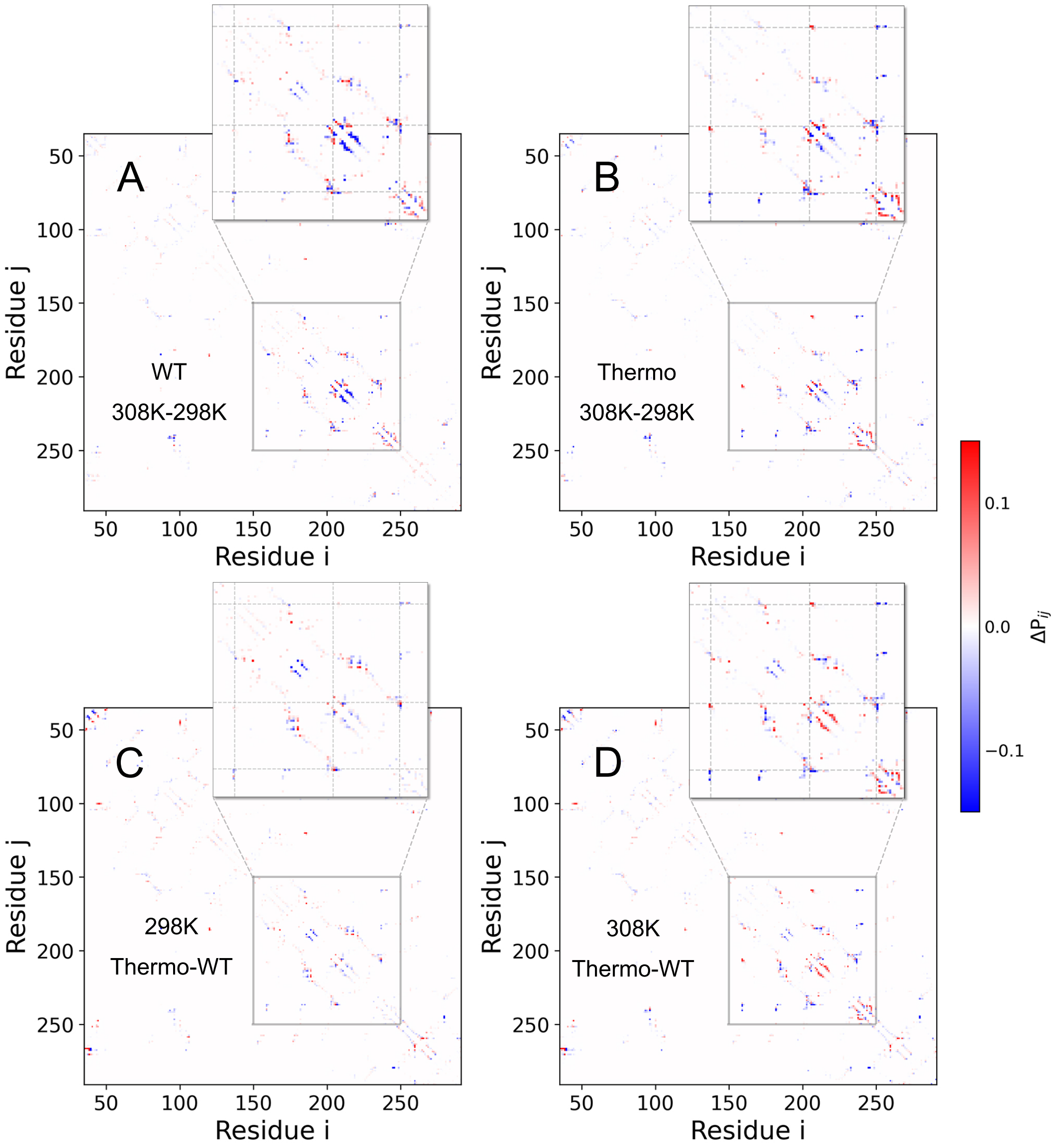


Figure S3: Differences in contact probability maps between WT- and Thermo-PETase at different temperatures. Contact probability difference between 298K and 308K of (A) WT- and (B) Thermo-PETase, respectively. Contact probability differences between WT- and Thermo-PETase at (C) 298K, and (D) 308K, respectively.

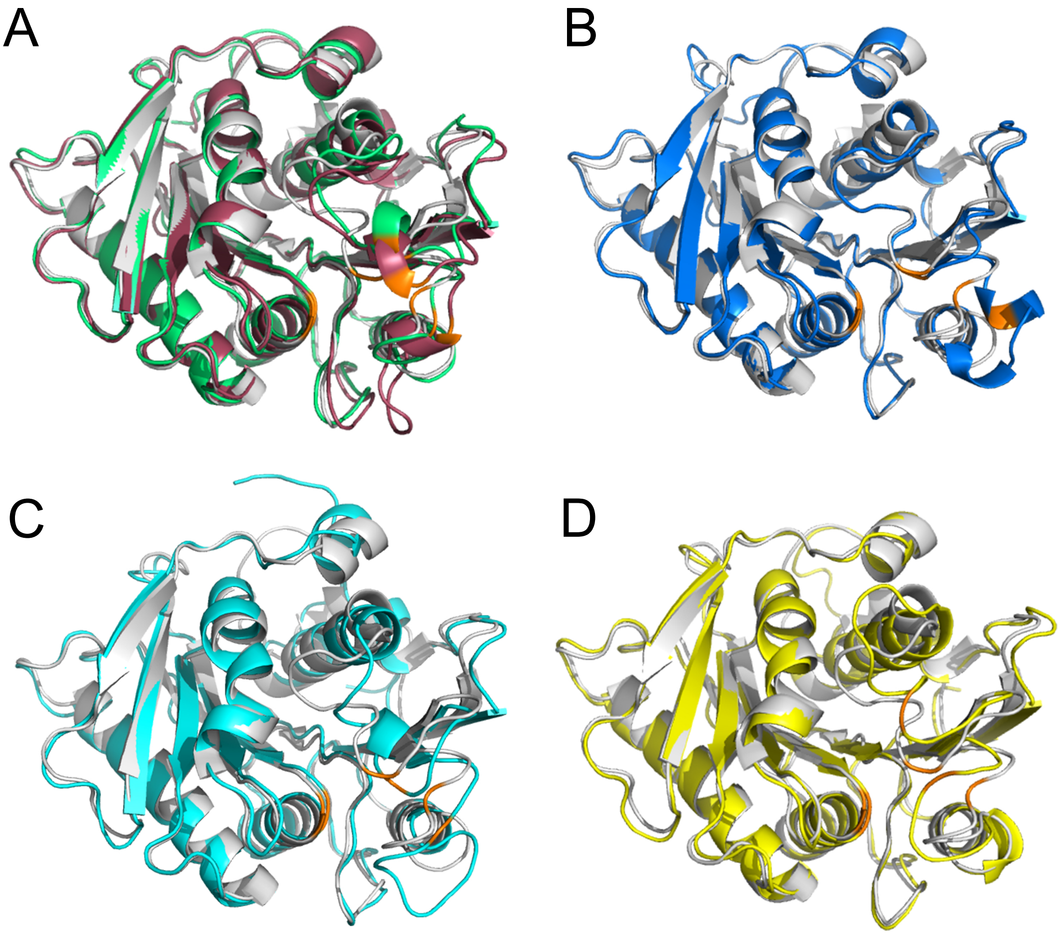


Figure S4: Global structural illustrations of PETase at the (meta)stable states identified on the FELs. The structures of PETase at the states are colored in the same scheme as they are in Figure 2. (A) State 3 (green) and state 4 (red). (B) State 5 (blue). (C) State 6 (cyan). (D) State 7 (yellow). State 1 in (A-D) is colored in grey. The three catalytic residues (Ser160, Asp206 and His237) are colored in orange.

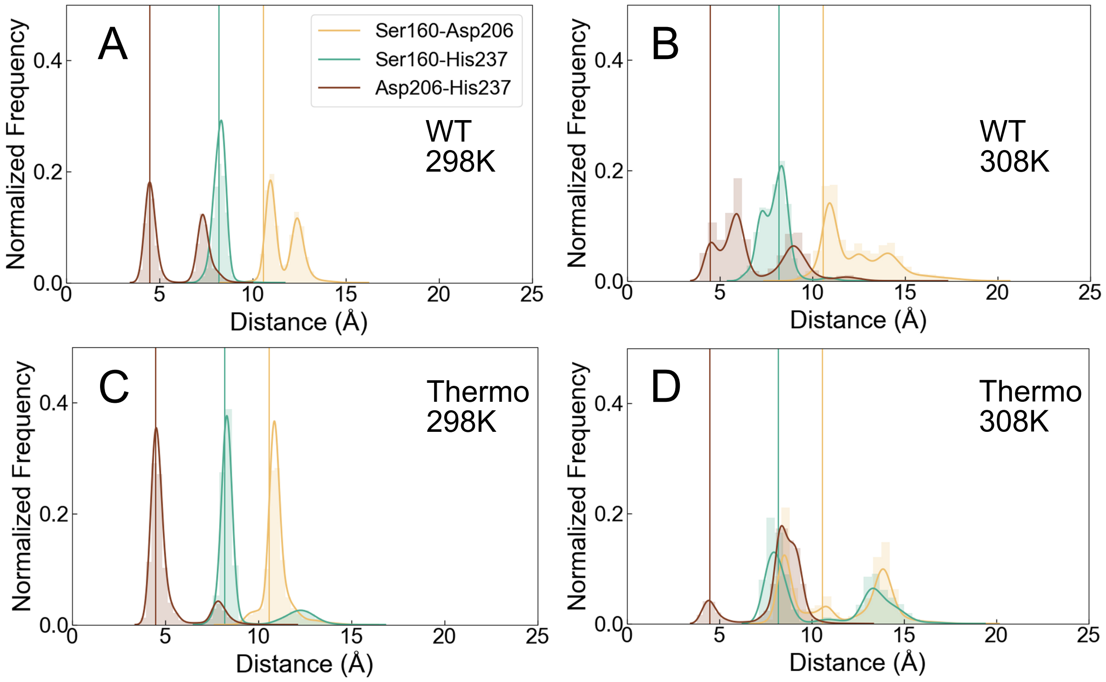


Figure S5: Pairwise distance distributions of the three residues at the catalytic site (Ser160, Asp206 and His237). (A) Distributions for WT-PETase at (A) 298K and (B) 308K. Distributions for Thermo-PETase at (C) 298K and (D) 308K.

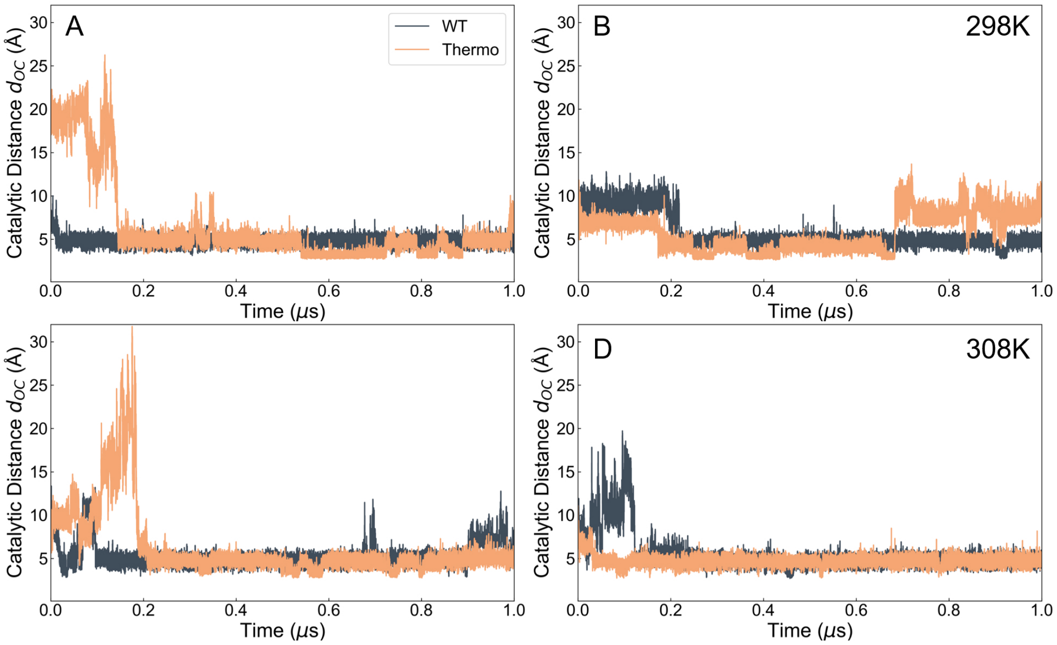


Figure S6: Evolutions of distance ($d_{OC}$) between the oxygen $O_\gamma$ of the catalytic Ser160 and the carbonyl carbon $C$ of the substrate during MD simulations for WT- and Thermo-PETase at 298K and 308K.

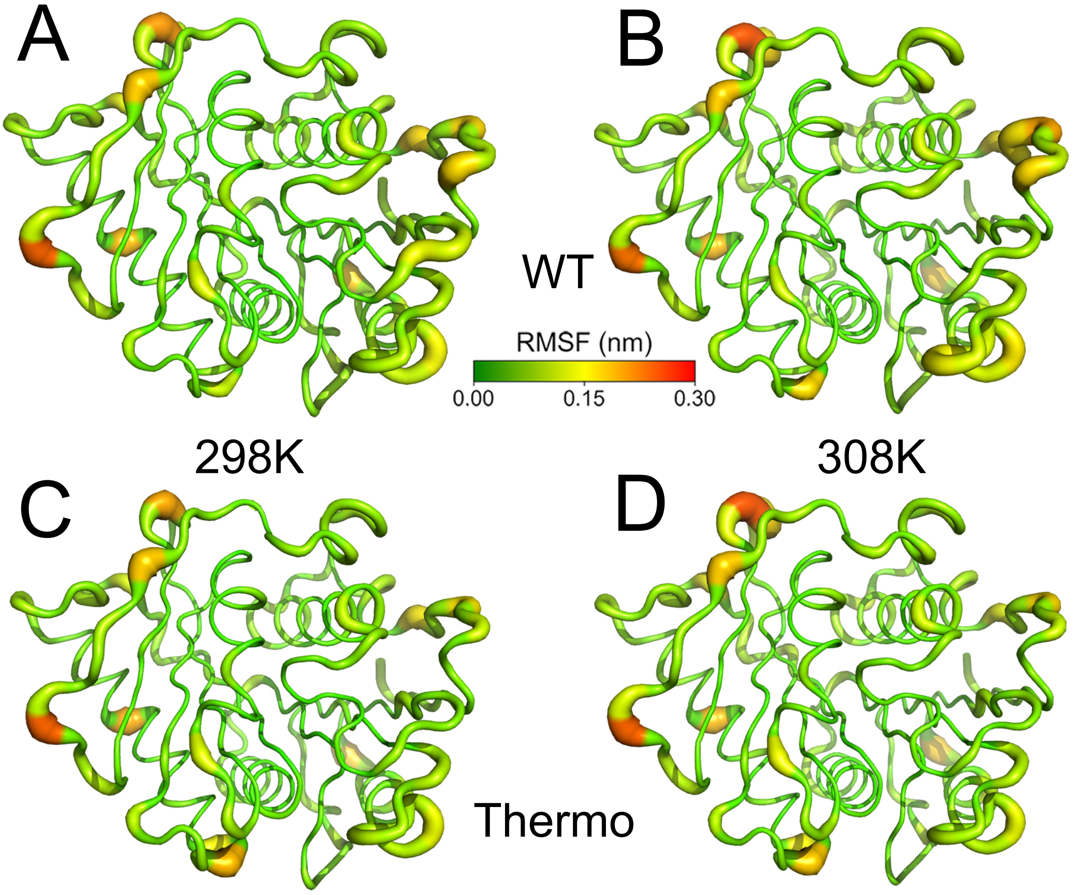


Figure S7: RMSF values at 298K and 308K for WT- and Thermo-PETase projected onto the corresponding native structures.

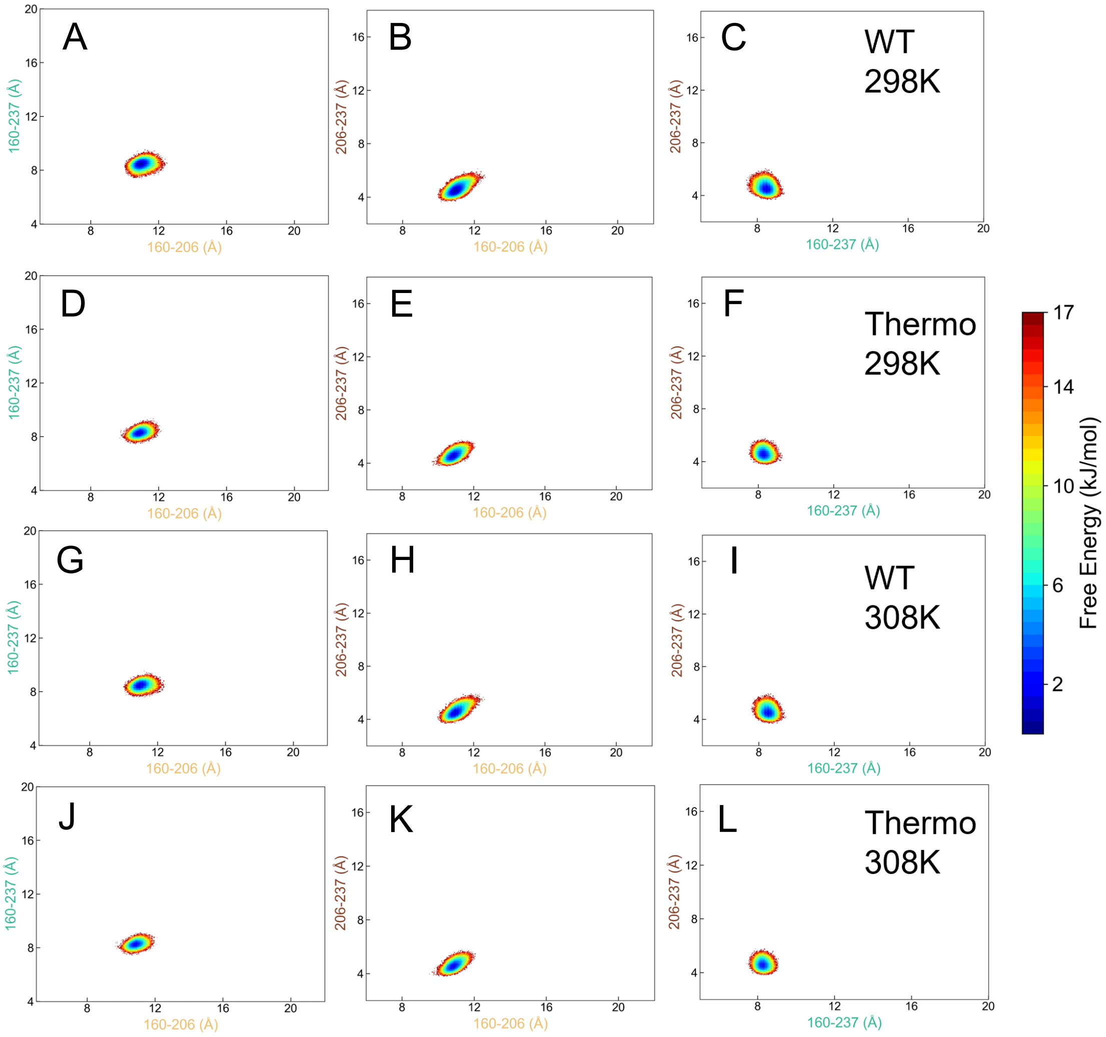


Figure S8: FELs projected onto the distances between every pair of the catalytic triad of Ser160, Asp206 and His237 after substrate binding.

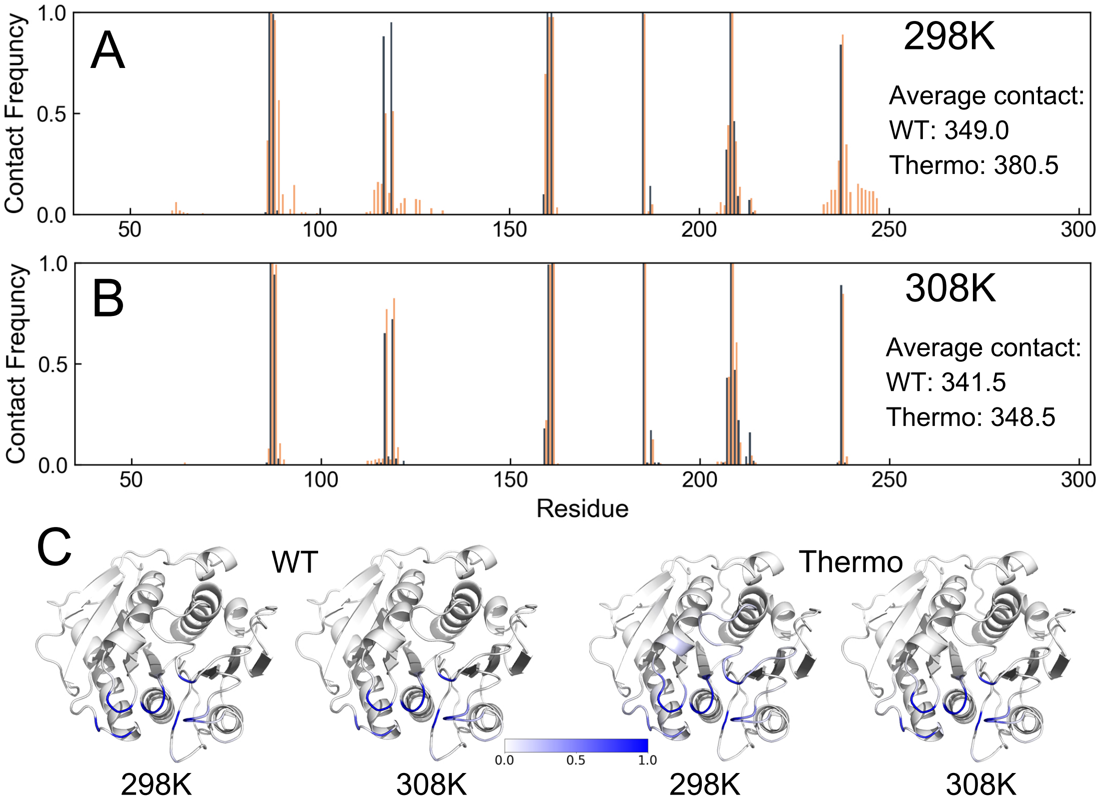


Figure S9: Contacts between PETase and 4PET. The contact frequency between PETase and PET along the residue for WT- and Thermo-PETase at (A) 298K and (B) 308K. (C) Contact frequencies in (A) and (B) projected onto the corresponding native structures of WT- and Thermo-PETase.

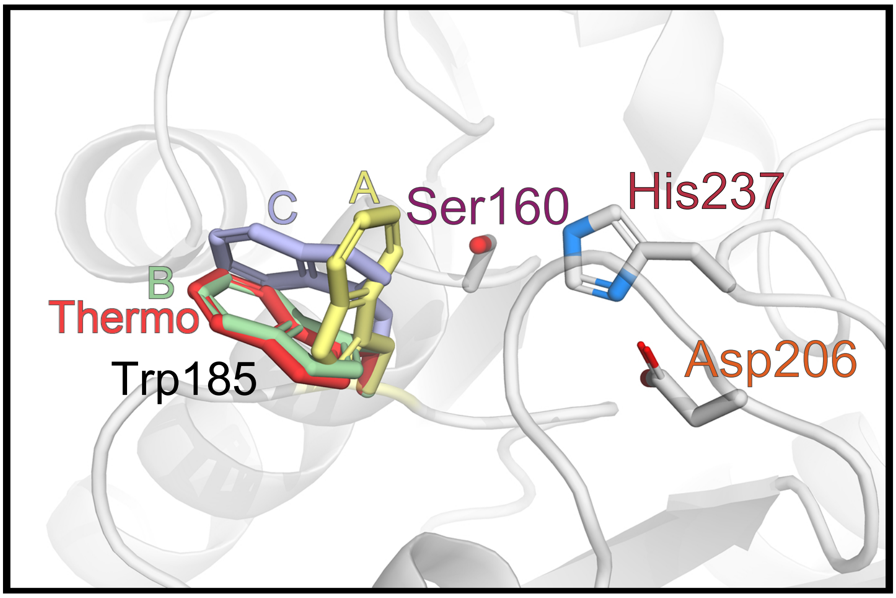


Figure S10: Multiple conformations of Trp185 in crystal structures of WT-PETase (PDB: 5XG0 [3]) and Thermo-PETase (PDB: 6IJ6 [4]). The side chain of Trp185 and catalytic residues are shown with sticks. The conformations of Trp185 in WT-PETase are colored in yellow ("conformer A"), green ("conformer B") and purple ("conformer C"). The conformation of Trp185 in Thermo-PETase is colored in red, similar to "type B" conformation. Noteworthy, the conformation of Trp185 in the presence of the substrate analogue HEMT adopts the "conformer B" conformation [3].

### References


[1] Garrett M. Morris, Ruth Huey, William Lindstrom, Michel F. Sanner, Richard K. Belew, David S. Goodsell, and Arthur J. Olson. Autodock4 and autodocktools4: Automated docking with selective receptor flexibility. *Journal of computational chemistry*, 30(16):2785–2791, December 2009.

[2] Jerome Eberhardt, Diogo Santos-Martins, Andreas F. Tillack, and Stefano Forli. Autodock vina 1.2.0: New docking methods, expanded force field, and python bindings. *Journal of Chemical Information and Modeling*, 61(8):3891–3898, August 2021.

[3] Xu Han, Weidong Liu, Jian-Wen Huang, Jiantao Ma, Yingying Zheng, Tzu-Ping Ko, Limin Xu, Ya-Shan Cheng, Chun-Chi Chen, and Rey-Ting Guo. Structural insight into catalytic mechanism of pet hydrolase. *Nature communications*, 8(1):2106, 2017.

[4] Hyeoncheol Francis Son, In Jin Cho, Seongjoon Joo, Hogyun Seo, Hye-Young Sagong, So Young Choi, Sang Yup Lee, and Kyung-Jin Kim. Rational protein engineering of thermo-stable petase from *ideonella sakaiensis* for highly efficient pet degradation. *ACS Catalysis*, 9(4):3519–3526, April 2019.
